# Analysing biodiversity observation data collected in continuous time: Should we use discrete- or continuous-time occupancy models?

**DOI:** 10.1101/2023.11.17.567350

**Authors:** Léa Pautrel, Sylvain Moulherat, Olivier Gimenez, Marie-Pierre Etienne

## Abstract

1. Biodiversity monitoring is undergoing a revolution, with fauna observations data being increasingly gathered continuously over extended periods, through sensors like camera traps and acoustic recorders, or via opportunistic observations. These data are often analysed with discrete-time ecological models, requiring the transformation of continuously collected data into arbitrarily chosen non-independent discrete time intervals. To overcome this issue, ecologists are increasingly turning to the existing continuous-time models in the literature. Closer to the real detection process, they are lesser known than discrete-time models, not always easily accessible, and can be more complex. Focusing on occupancy models, a type of species distribution models, we asked ourselves: Should we dedicate time and effort to learning and using these continuous-time models, or can we go on using discrete-time models?
2. We conducted a comparative simulation study using data generated within a continuous-time framework. We assessed the performance of five static occupancy models with varying detection processes: discrete detection/non-detection process, discrete count process, continuous-time Poisson process, and two types of modulated Poisson processes. Our goal was to assess their abilities to estimate occupancy probability with continuously collected data. We applied all models to empirical lynx data as an illustrative example.
3. In scenarios with easily detectable animals, we found that all models accurately estimated occupancy. All models reached their limits with highly elusive animals. Variation in discretisation intervals had minimal impact on the discrete models’ capacity to estimate occupancy accurately.
4. Our study underscores that opting for continuous-time models with an increased number of parameters, aiming to get closer to the sensor detection process, may not offer substantial advantages over simpler models when the sole aim is to accurately estimate occupancy. Model choice can thus be driven by practical considerations such as data availability or implementation time. However, occupancy models can encompass goals beyond estimating occupancy probability. Continuous-time models, particularly those considering temporal variations in detection, can offer valuable insights into specific species behaviour and broader ecological inquiries. We hope that our findings offer valuable guidance for researchers and practitioners working with continuously collected data in wildlife monitoring and modelling.

## 1 Introduction

The alarming decline of biodiversity has led to a scientific, ethical, and legal need to better understand its drivers in order to protect nature more effectively (IPBES, 2019). With the reinforcement of regulations and recommendations for achieving the objectives of no net loss of biodiversity, the need for wildlife monitoring is growing rapidly (UNECE, 2023). Concurrently, the development of increasingly sophisticated and accessible technologies is leading to a digital revolution. Sensors, such as camera traps or autonomous recording units, are now available to address current ecological challenges (Burton et al., 2015; Potamitis et al., 2014).

Sensors offer many advantages compared to traditional field observations by naturalists. They are non-invasive, often cost-effective, particularly adapted to observe some elusive or shy species, potentially in challenging terrain, and they can improve reproducibility and protocol standardisation (Steenweg et al., 2017; Zwerts et al., 2021). Sensors are therefore good candidates for setting up large-scale monitoring (Oliver et al., 2023) and collaborations such as Biodiversity Observation Networks (Gonzalez et al., 2023). Policies now emphasise the use of sensors, big data and artificial intelligence to improve knowledge and understanding of species and ecosystems, such as the International Union for Conservation of Nature (IUCN) Nature 2030 programme (IUCN, 2021) or the Biodiversa+ European Biodiversity Partnership (Høye et al., 2022; Vihervaara et al., 2023).

We often use ecological models to analyse observation data for monitoring purposes. These models typically assess the presence (Guillera-Arroita, 2017) or abundance (Gilbert et al., 2021) of a species, often while considering the relation with environmental factors. They can be used for a particular species or within a multi-species framework (Pollock et al., 2014). These models produce actionable knowledge about species, influencing our actions and our approach to biodiversity conservation. For example, the area of occupancy, *i*.*e*. the spatial distribution where a species is present, is one of the criterion used by the IUCN to establish the Red list of Ecosystems (Rodríguez et al., 2015).

In this paper, we focus on occupancy models, a category of ecological models aiming to estimate species presence. Occupancy models, as introduced by MacKenzie et al. (2002), are hierarchical models that include two sub-models. The first sub-model describes the ecological process, occupancy, typically of interest to ecologists. The second sub-model accounts for measurement errors arising from imperfect detection. A site is said occupied when at least one individual went through it (Emmet et al., 2021). At a broader scale, occupancy corresponds to the proportion of sites within a study area that are occupied by the species (MacKenzie et al., 2002). The occupancy model proposed by MacKenzie et al. (2002) uses binary data (0 if the species was not detected, 1 if it was) at each site during each sampling occasion. This model has underpinned numerous occupancy studies in the last two decades, and was refined or adapted by many modelers (Bailey et al., 2014). These adaptations have given rise to new occupancy models, most of them aiming to mirror more closely the expected ecological or detection conditions, impacting the input data required by each model. We here focus on static occupancy models, in which the occupancy state of a site is assumed constant, without extinction-colonisation processes, as opposed to dynamic occupancy models.

Ecological models, including occupancy models, have historically been developed to analyse observation data collected by field operators during one or several short sampling occasions (Bailey et al., 2014). However, the deployment of sensors involves continuous data collection, often over long time periods (*e*.*g*. Cove et al., 2021; Cusack et al., 2015; Moore et al., 2020). For instance, Kays et al. (2020) recommend deploying sensors for three to five weeks at multiple locations to estimate relative abundance, occupancy, or species richness. Short-term deployments can equate traditional discrete sampling occasions. However, when sensors are stationed at the same location for extended periods, data is often discretised in order to use traditional models in discrete time. We suggest using the term **session** for these discretised time intervals, because they differ from traditional sampling occasions in two respects: *(1)* sampling occasions are determined before the data collection, whereas the discretisation is done after the data has been collected; and *(2)* sessions occur consecutively without any gaps between them, while the traditional sampling occasions are separated by periods of time when the site is not monitored.

Occupancy discrete-time models have been around for 20 years and are commonly used because they are relatively simple to implement. However, continuous-time ecological modelling is not new. The fist mention of a continuous-time model in the capture-recapture literature dates back to Becker (1984). It was not until the advent of sensors, which highlighted the limitations of discrete-time models, that modelers began to turn towards continuous-time models (Kellner et al., 2022; Rushing, 2023; Schofield et al., 2018). Nonetheless, continuous-time models are not a universal cure-all. Each family of models have their pros and cons.

### Discretisation simplifies the information

Discretisation is, in other words, an aggregation of data into sessions. This aggregation simplifies the data and blurs the residual variability, which can help in interpreting broad observed trends. But simplification involves information loss. Because accurately estimating occupancy relies on precisely assessing imperfect detection (Kellner & Swihart, 2014; Kéry & Schmidt, 2008), having more information about detection patterns could provide valuable insights, helping to disentangle the observation process from the ecological process of interest. This may lead to more accurate estimations of static occupancy. Dynamic occupancy models could represent the occupancy state in continuous time to go even further, potentially revealing fine patterns with ecological significance.

### Discretisation is arbitrary

Researchers usually choose the aggregation period so that the detection probability is not too low, and the occupancy probability is not estimated at its boundaries (close to 0 or 1). Schofield et al. (2018) highlighted that the chosen session length can impact abundance estimates with discrete-time capture-recapture models. Hence, it most likely impacts occupancy models outputs, as capture-recapture and occupancy models are very similar (the individual capture history equates the site “detection history”, MacKenzie et al., 2002). Eliminating arbitrary discretisation in occupancy modelling can enhance the method objectivity and reproducibility, and is expected to improve result reliability, at least compared to a non-optimal discretisation.

### Model complexity and data availability

Models with a continuous-time detection process are likely to overcome the limitations mentioned above. Furthermore, harnessing the richness of continuous-time data, researchers can customise models to replicate species-specific observation processes, providing valuable insights into animal behaviour and activity patterns (e.g. Distiller et al., 2020, with continuous-time spatial capture-recapture models). However, the potential drawback is the complexity of such models, that may render them less adapted to derive ecological insights such as occupancy from small data sets. Additionally, if the system is not assumed to be constant over time, continuous-time covariates are necessary for a continuous-time model, and these covariates are often not readily available.

### Importance of discretisation versus distribution law

In response to Schofield et al. (2018), Zhang and Bonner (2020) demonstrated that differences in inference with capture-recapture discrete-time models were not attributed to varying data discretisation scales but rather to the choice of the distribution law for modelling the detection process. When dealing with mathematically equivalent models, both continuous- and discrete-time models would yield equivalent outcomes. Consequently, the decision between discrete- and continuous-time models is less significant and impactful than choosing a model with a different distribution. We also note that the choice of distribution law to model detection influences the model parameters. For instance, we find detection rates more practical than detection probabilities, improving the comparability of studies as probabilities are only meaningful at the scale of the discretised period.

Users select an occupancy model depending on a trade-off between model performance and implementation cost. This cost encompasses factors such as model familiarity, programming if necessary, and accessibility to data, all of which can be influenced by the complexity of the model. Existing comparisons between discrete- and continuous-time models are presented in papers introducing new continuous models, focusing on evaluating the new model formulation, and often limited to just two models (Emmet et al., 2021; Guillera-Arroita et al., 2011). Other studies compare models that use time-to-first-detection data with those using repeated measures to improve field survey methods conducted by human observers (Halstead et al., 2021; Henry et al., 2020; Priyadarshani et al., 2022).

In this paper, we investigate whether continuous-time modelling is beneficial for occupancy estimation using sensor-based observation data and under which circumstances. We conduct a comprehensive comparison of five occupancy models, varying in the complexity of their detection processes. These five models cover a large scope of single-species static occupancy models with no false positives (MacKenzie et al., 2004). We omitted time-to-first-detection models. Although often appropriate to analyse data from time-optimised human-based surveys (Halstead et al., 2021; Henry et al., 2020; Priyadarshani et al., 2022; Priyadarshani et al., 2024), using only the first detection from sensor-based data amounts to discarding lots of informative data. Therefore, we considered only time-to-each-detection for continuous-time models. We also omitted models that consider abundance-induced detection heterogeneity, like the Royle-Nichols model (Royle & Nichols, 2003).

We compare the ability of occupancy models to retrieve the simulated occupancy probability using complementary comparison metrics, measuring accuracy, bias, and precision (Liemohn et al., 2021). To fully control the environment, we simulate continuous detection data. This allows us to explore how the rarity and elusiveness of the target species influences the model’s ability to retrieve the occupancy. We also simulate extreme cases to refine the models’ application limits. As an illustrative example, we used the five compared models to analyse continuously collected empirical lynx (*Lynx lynx*) data observed through camera traps. We aim to offer recommendations for choosing discrete- or continuous-time models and to discuss various considerations that researchers should address when analysing fauna observation data collected through sensors.

## 2 Material and methods

### 2.1 Occupancy models

In this section, we describe the five hierarchical occupancy models compared, with an ecological process modelling presence or occupancy, and an observation process addressing imperfect detection. The occupancy sub-model is consistent across all five models, while the detection sub-model differs. Fig. 1 provides an overview of the formulation and input data of the considered models, which are described in detail in the following paragraphs. The mathematical notation is listed in Table 1.

**Table 1:**
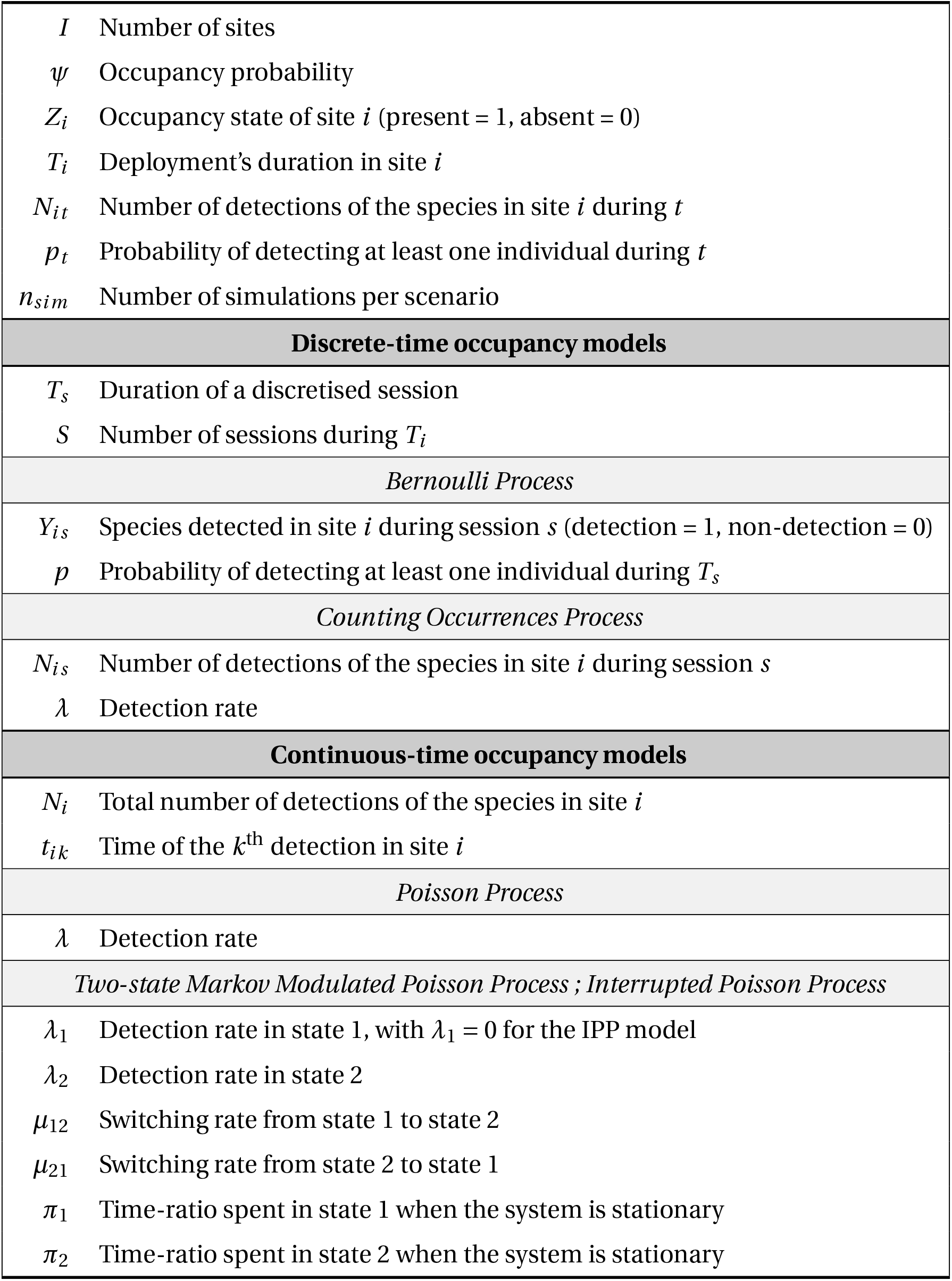
Notation.

**Figure 1:**
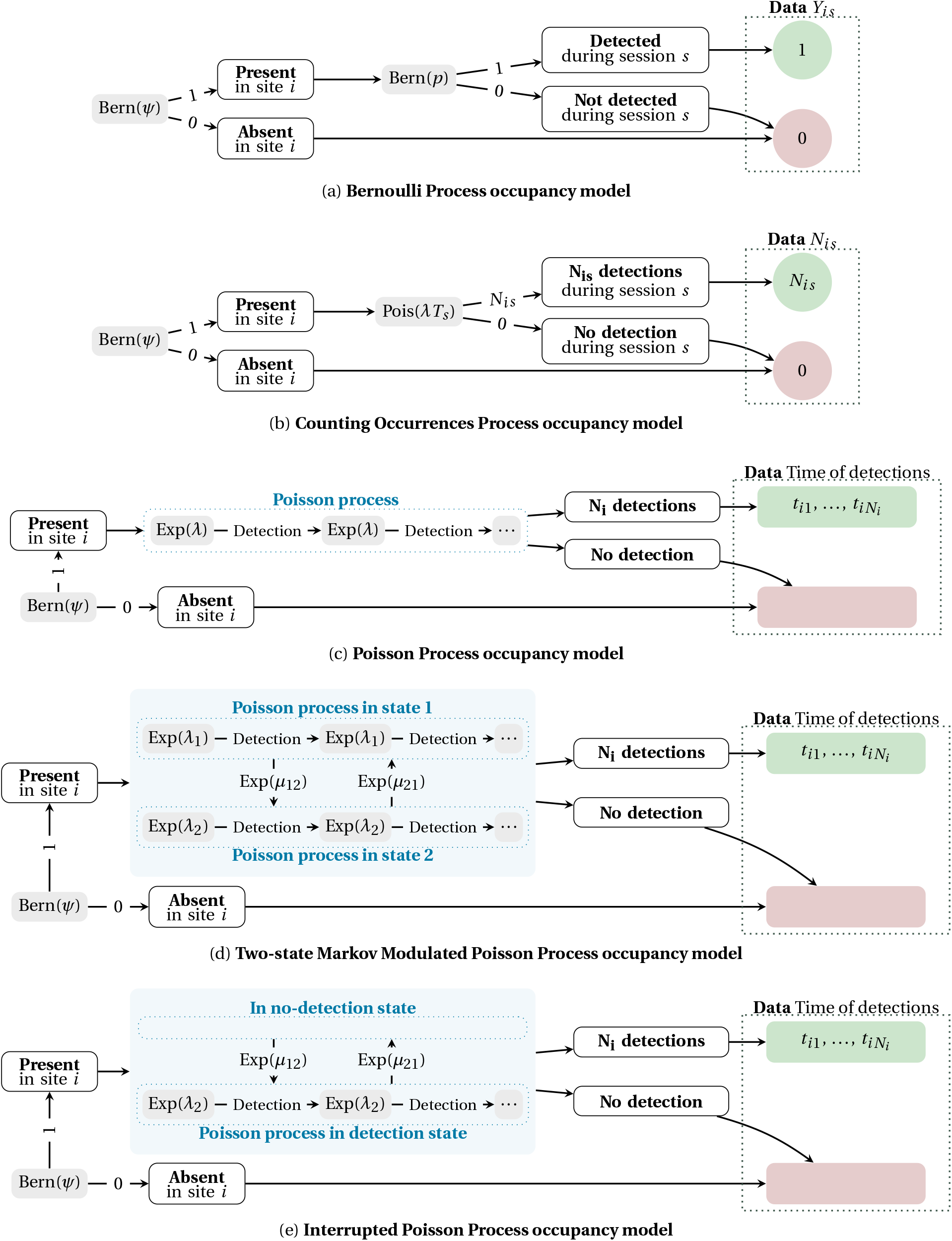
Five occupancy models compared. With: *ψ* the occupancy probability; **(a) BP** *p* the detection probability; *Y*_*is*_ the detection/non detection observed in site *i* during session *s*; **(b) COP** *λ* the detection rate; *T*_*s*_ the duration of a session; *N*_*is*_ the number of detections in site *i* during session *s*; **(c) PP** *λ* the detection rate; *N*_*i*_ the number of detections in site *i* ; *t*_*ik*_ the time of the *k*^th^ detection in site *i* ; **(d) 2-MMPP** and **(e) IPP** *λ*_1_ the detection rate in state 1; *λ*_2_ the detection rate in state 2; *μ*_12_ the switching rate from state 1 to state 2; *μ*_21_ the switching rate from state 2 to state 1; *N*_*i*_ the number of detections in site *i* ; *t*_*ik*_ the time of the *k*^th^ detection in site *i*.

#### 2.1.1 Occupancy sub-model

Across all five models, the occupancy sub-model is identical, assuming that the occupancy state of site *i, Z*_*i*_, follows a Bernoulli distribution with parameter *ψ*, the occupancy probability. The sites are assumed independent, regarding both occupancy and detection.

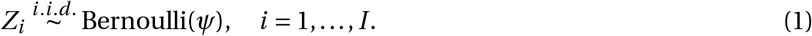

If the species is detected at least once in a site, that site is considered occupied, assuming no false positives. Temporal changes in occupancy are not considered; for simplicity, we focus on single-season occupancy models with no covariates.

#### 2.1.2 Detection sub-model

Two models rely on the time discretisation of the sensor-based observation data (Bernoulli Process (BP) and Counting Occurrences Process (COP)), while three others consider the detection as the realisation of a continuous-time stochastic process (Poisson Process (PP), Two-state Markov Modulated Poisson Process (2-MMPP) and Interrupted Poisson Process (IPP)). Their growing complexity, associated with an expected closer alignment with reality, influences the input data required for each model. Our primary focus is to determine if more complex representations of the detection process lead to improved estimates of occupancy probability, with minimised error and bias.

##### Bernoulli Process (BP)

In the classical occupancy model proposed by MacKenzie et al. (2002), the raw data are aggregated and simplified. The continuous data are aggregated into *S* sessions of duration *T*_*s*_, and simplified into the observation *Y*_*i s*_, which is 1 if at least one detection occurs during session *s* at site *i*, and 0 otherwise. Conditionally on the occupancy state *Z*_*i*_ of site *i*, the model assumes that the distribution of the variable of interest *Y* depends on *p* the probability of detecting at least one individual during a session:

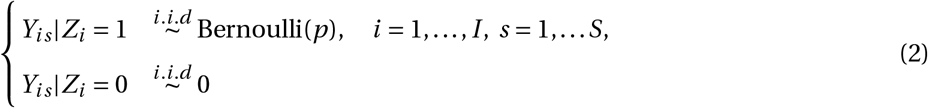

##### Counting Occurrences Process (COP)

In the BP model, detecting few or many individuals during a session leads to the same observation *Y*_*i s*_ *=* 1, although it corresponds to very different situations. We simplified the approach proposed by Emmet et al. (2021) to avoid references to secondary sessions and to use probability. As a result, its likelihood has been adjusted and is provided in supplementary information.

Although the data is aggregated by session like in the BP model, more information is retained since this approach models *N*_*i s*_, the number of individuals seen at site *i* during session *s*. Conditionally on the occupancy state *Z*_*i*_ of site *i*, as it is typical for count data, the COP model assumes that the number of detections *N*_*i s*_ follows a Poisson distribution of parameter *λ* the detection rate multiplied by *T*_*s*_ the session duration:

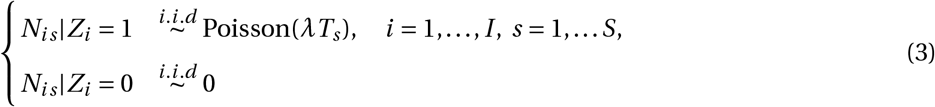

In practical terms, if the time-unit is a day, then when the detection rate *λ =* 3, there are on average three individuals detected by day. If each session lasts a week, *T*_*s*_ *=* 7, then there are on average *λT*_*s*_ *=* 3 *×* 7 *=* 21 individuals detected per session. The probability of detecting *k* individuals during a session is 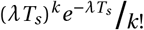. With this example, in an occupied site during a session, there is a 8.67% chance of detecting 21 individuals, a 0.35% chance for 10 individuals, and a 7.58*e*^*−*8^% chance of detecting nothing.

##### Poisson Process (PP)

Unlike the two previous models which required data discretisation, the PP occupancy model proposed by Guillera-Arroita et al. (2011) uses the time of detections as data, with *t*_*i j*_ the time of the *j* ^th^ detection in site *i*. These raw, unaggregated data retain all of its information. The time of detections are transformed into interdetection times to calculate the likelihood of these data given the model and its parameters. The first interdetection time is usually defined as the time between the deployment beginning and the first detection, the second as the time between the first detection and the second, and so forth. The last value in this vector can be defined as the time between the last detection and the end of deployment. If the time at which the deployment ended is not known, *e*.*g*. because the battery died, the likelihood can be adapted so that this last value can be the time between the second-to-last detection and the last detection (Guillera-Arroita et al., 2011).

When the site *i* is occupied, the detection process is modeled as a homogeneous Poisson point process of parameter *λ*, the detection rate. This means that the interdetection times are exponential variables with rate *λ*. In practical terms, if the time-unit is a day, then a detection rate *λ =* 3 means that on average, three individuals are seen per day. The average time between two detections is ⅓ of a day.

One property of a Poisson process of parameter *λ* is that the number of detections over a period of time *T* follows a Poisson distribution with parameter *λT*. This model is therefore mathematically equivalent to the COP model presented above (Zhang & Bonner, 2020). Nonetheless, using the raw data could enable ecologists to delve deeper and consider the detection rate heterogeneity with the model residuals.

##### Two-state Markov Modulated Poisson Process (2-MMPP)

The 2-MMPP occupancy model was also proposed by Guillera-Arroita et al. (2011) and uses the time of detection events as data, transformed into interdetection times. Unlike the PP occupancy model, which assumes that detection events happen at an homogeneous rate, this model incorporates some hetereogeneity. This approach is likely more representative for many species, considering various ecological processes that can lead to temporal clustering in detection events. Examples include seasonal activity patterns, or, at finer temporal scales, daily activity patterns, and interactions with other species, among others. In the 2-MMPP occupancy model, when the site *i* is occupied, the detection process is modeled as a system of Poisson processes with two different rates. When the system is in state 1, the detection events are modeled by a Poisson process of parameter *λ*_1_. In state 2, the rate is *λ*_2_. This is a two-state continuous-time Markov chain, where the system switches from one hidden state to the other, with parameters *μ*_12_ (switching rate from state 1 to state 2) and *μ*_21_ (switching rate from state 2 to state 1).

With day as the time-unit and a set of parameters of *λ*_1_ *=* 1, *λ*_2_ *=* 5, *μ*_12_ *=* ^1^/_15_, *μ*_21_ *=* 1, this means that:

- State 1 is a low-detection state with 1 detection per day on average (*λ*_1_), State 2 is a high-detection state with 5 detections per day on average (*λ*_2_)
- When the system is in state 1, there is ^1^/_15_ switch to state 2 per day on average (*μ*_12_), corresponding to 15 days spent on average in state 1 before switching to state 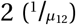. When the system is in state 2, there is 1 switch to state 1 per day on average (*μ*_21_), corresponding to 1 days spent on average in state 2 before switching to state 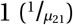
- The system is in state 1 for 93.75% of the deployment time on average (*π*_1_ in Equation 4), and in state 2 for 6.25% of the time (*π*_2_ in Equation 4)
- In an occupied site, there are on average 1.25 detections per day (Equation 5) and the variance of the number of daily detections is 4.11 (Equation 6)

The proportion of time spent in each state when the system is stationary is the steady-state vector Π of the Markov chain for a 2-MMPP, is presented in Equation 4 (Fischer & Meier-Hellstern, 1993).

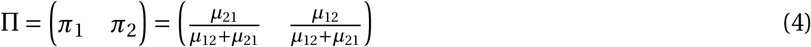

The number of events (here *N*_*i t*_ the number of detections at site *i* taking place during an observation time *t*) of a 2-MMPP is described by its expected value 𝔼 [*N*_*i t*_ ] in Equation 5 and by its variance 𝕧 [*N*_*i t*_ ] in Equation 6 (see Supplementary Informations and Bhat, 1992).

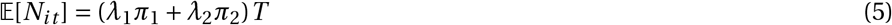

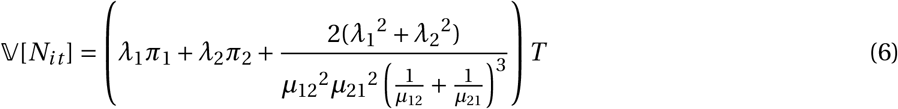

The probability of having at least one detection during an observation period of duration *T*, written *p*_*t*_, is given in Equation 7, with *exp* the matrix exponential function (from Guillera-Arroita et al., 2011, section 4.2).

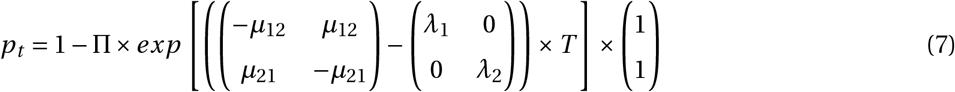

MMPPs are a type of Cox processes (Cox, 1955). 2-MMPPs can also be referred to as switched Poisson processes (SPP, Arvidsson and Harris, 1991) or as a doubly stochastic Poisson processes (Bhat, 1992, 1994). For simplicity, we focused on two-state models in this comparison. However, in specific ecological contexts, considering more states may be relevant. For more informations on MMPPs in general, with possibly more than 2 states, see Fischer and Meier-Hellstern (1993), Guillera-Arroita (2012), and Rydén (1994).

##### Interrupted Poisson Process (IPP)

The IPP occupancy model is a special case of a 2-MMPP where there are no detections in one of the two states. This modelling approach is intuitive for ecological settings where we expect periods without detection, such as diurnal species (active and observed during the day, inactive thus unobserved at night) or gregarious species (extended periods with no detections, and at the passage of a herd, shorter periods with numerous detection events). In such contexts, the IPP model, more parcimonious with one less parameter, could be more adapted than the 2-MMPP model, which might still estimate a near-zero detection rate and produce equivalent results. Since usually, *λ*_1_ *< λ*_2_ (Skaug, 2006), an IPP is a 2-MMPP with *λ*_1_ *=* 0 constrained.

### 2.2 Simulation study

#### 2.2.1 Continuous detection data simulation

We simulated detection data in *I =* 100 sites, with one deployment per site of *T*_*i*_ *=* 100 time-units. For the sake of simplicity, one time-unit corresponds to one day throughout this article. We simulated data with various occupancy probability and detection parameters. All simulation parameters are described in Table 2. In detection scenarios (a) and (b), we simulated extreme cases of species elusiveness to identify the models’ limits and behaviour in extreme situations, even if we expect these to produce insufficient data to perform occupancy modelling. We carried out *n*_*sim*_ *=* 500 simulations per simulation scenario.

**Table 2:** Simulation parameters. With *p*_100_ the probability of having at least one detection during a deployment of *T*_*i*_ *=* 100 days at an occupied site (Equation 7); *p*_1_ the probability of having at least one detection during 1 day (Equation 7); 𝔼 [*N*_100_] the expected number of detections during a deployment of *T*_*i*_ *=* 100 days at an occupied site (Equation 5); 𝕧 [*N*_100_] the variance of the number of detections during a deployment of *T*_*i*_ *=* 100 days at an occupied site (Equation 6)

| (a) General parameters |  | (b) Parameters of the seven detection scenarios |  |  |  |  |  |  |  |  |
| --- | --- | --- | --- | --- | --- | --- | --- | --- | --- | --- |
| $I$ | 100 sites | | $\lambda_1$ | $\lambda_2$ | $\mu_{12}$ | $\mu_{21}$ | $p_{100}$ | $p_1$ | $\mathbb{E}[N_{100}]$ | $\mathbb{V}[N_{100}]$ |
| $T_i$ | 100 days | (a) | 0.00 | 1.00 | $1/15$ | 24 | 0.23 | 0.003 | 0.28 | 0.30 |
| $n_{sim}$ | 500 simulations per scenario | (b) | 0.00 | 5.00 | $1/15$ | 24 | 0.68 | 0.01 | 1.39 | 1.96 |
| $\psi$ | 0.10, 0.25, 0.50, 0.75, 0.90 | (c) | 0.00 | 1.00 | $1/15$ | 1 | 0.96 | 0.04 | 6.25 | 17.24 |
| $T_s$ | 30 (month), 7 (week), 1 (day) | (d) | 0.25 | 0.25 | $1/15$ | $1/10$ | 1.00 | 0.22 | 25.00 | 61.00 |
| | | (e) | 0.00 | 5.00 | $1/15$ | 1 | 1.00 | 0.09 | 31.26 | 306.03 |
| | | (f) | 0.00 | 1.00 | $1/15$ | $1/10$ | 1.00 | 0.26 | 40.01 | 327.98 |
| | | (g) | 0.00 | 5.00 | $1/15$ | $1/10$ | 1.00 | 0.42 | 200.06 | 7399.34 |

The occupancy status of each site was determined as the outcome of a Bernoulli trial with probability *ψ*. The detection process was simulated as a 2-MMPP of parameters *λ*_1_, *λ*_2_, *μ*_12_, *μ*_21_, using R version 4.2.3 (R Core Team, 2023). The state at the beginning of a deployment was drawn according to the stationary distribution, as a random sampling with probability *π*_1_ (*resp. π*_2_) of being in state 1 (*resp. 2*). Until the end of the deployment, the time to next event was a draw from an exponential distribution with parameter *μ*_12_ *+λ*_1_ in state 1, and with parameter in state 2. In state 1, this event was either a detection with probability 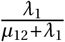, or a switch to state 2. In state 2, it was either a detection with probability 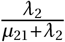, or a switch to state 1 (Fig. 2).

**Figure 2:**
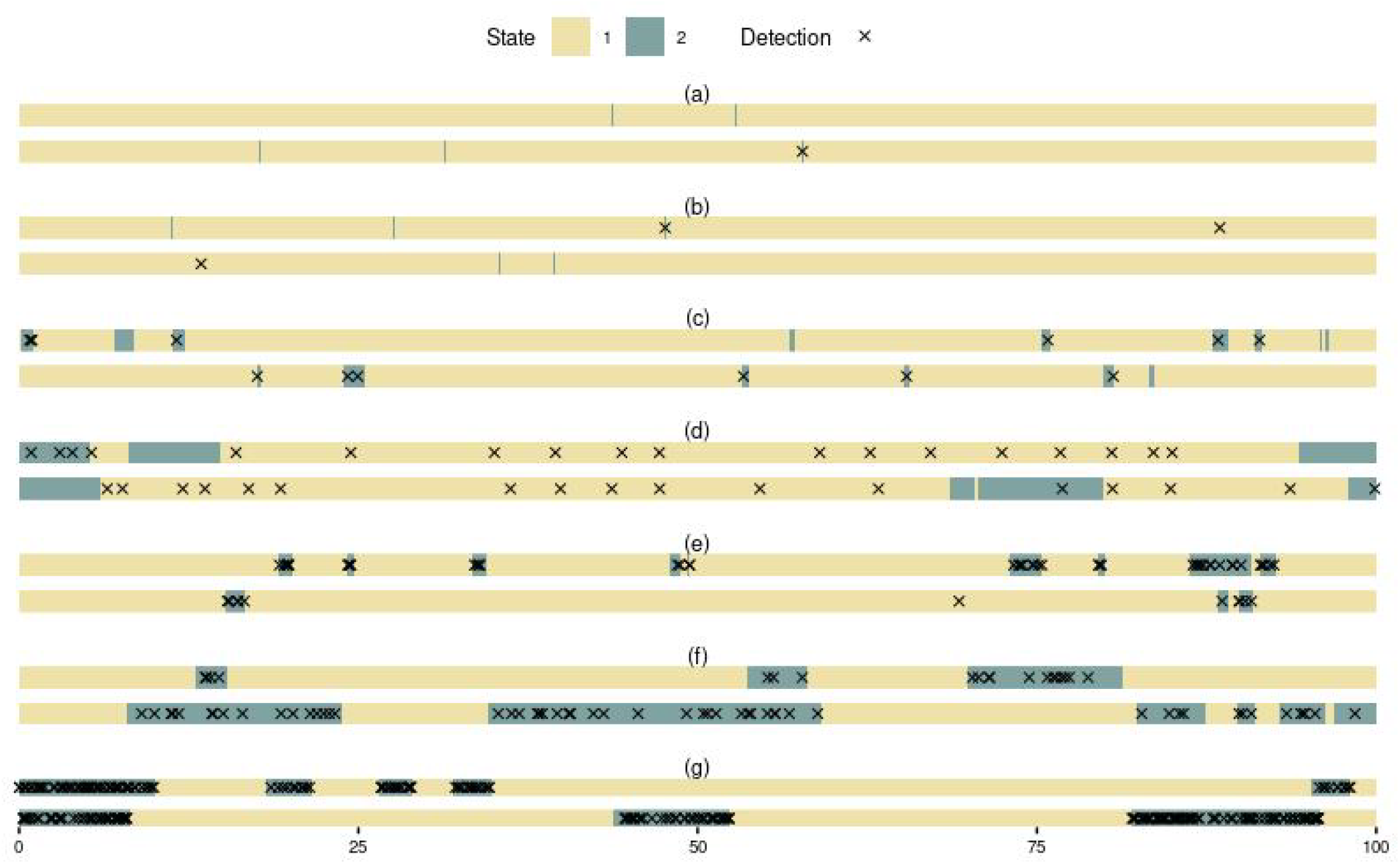
Simulated detection data. To help understand the impact of the detection parameters, two examples are given per detection scenario. With scenarios (a to g) described in Table 2. The detection process is simulated in an occupied site during 100 days.

##### Discretisation into sessions

For the two models that required discretisation into sessions, we used three levels of discretisation: monthly, weekly, and daily. Incomplete sessions are deemed invalid and will be excluded from the analysis. Consequently, when the data is discretised into months, there are three sessions consisting of 30 days each, and the detection data of the last 10 days of each deployment is disregarded. Similarly, when the data is discretised into weeks, there are 14 sessions of 7 days each, the last 2 days of each deployment is discarded.

#### 2.2.2 Frequentist parameter estimation

We estimated models parameters by maximum likelihood estimation and implemented it in R version 4.2.3 (R Core Team, 2023). For the COP, PP, 2-MMPP and IPP models, we used the `optim` function from the `stats` package (R Core Team, 2023) to maximise the log-likelihood. For the BP model, we used the function `occu` from the `unmarked` package version 1.3.2 (Fiske & Chandler, 2011), which calls the same `optim` function. We used the `Nelder-Mead` algorithm to maximise the likelihood. To reduce the optimisation time, we used the simulated parameters as the initial parameters to start the optimisation algorithm. The likelihood maximisation methodology was equivalent for the 5 models, making their results comparable. In order to perform unconstrained optimisation, we applied a logit transformation to the probabilities (*ψ, p*) and a log transformation to rates (*λ, λ*_1_, *λ*_2_, *μ*_12_ and *μ*_21_). In addition, we fitted the models with the BFGS optimisation algorithm. The results are not shown here but presented in supplementary information.

#### 2.2.3 Performance comparison for occupancy probability estimation

For each simulation scenario, we calculated the Root Mean Square Error (RMSE, Equation 8) as an error metric, measuring the absolute difference between the models’ point estimates of occupancy probability 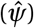 and the ground-truth occupancy probability (*ψ*), used to simulate data sets of this simulation scenario.

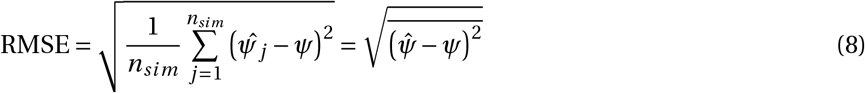

To complete this metric, we calculated absolute bias (AB, Equation 9) to better understand if this error was due to under-estimation or over-estimation of *ψ*.

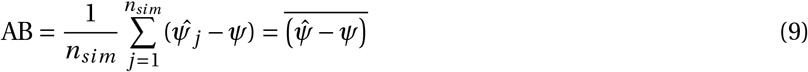

To compare the distributions of the point estimates 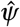 of the five different models and the different discretisations for BP and COP, we performed a Kruskal-Wallis test for each simulated scenario. We also conducted Wilcoxon tests with Bonferroni correction and visualised the distribution of 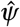.

We calculated for each inference the 95% confidence interval (CI) of the occupancy probability. To summarise this information for all the *n*_*sim*_ simulations by model in each simulation scenario, we used two metrics, the coverage (Equation 10) and the average range of the confidence interval (ARCI, Equation 11). We note *CI*_*l*_ and *CI*_*u*_ the lower and upper bounds of the 95% confidence interval of the estimated occupancy probability.

Coverage is the proportion of simulations for which the true simulated occupancy probability (*ψ*) is within the 95% CI of the estimated occupancy probability. In other words, coverage can be interpreted as the percentage of good predictions of the occupancy probability by a model.

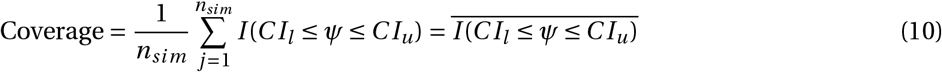

The average range of the 95% confidence interval measures the precision of the estimation, with the width of the confidence interval. It completes coverage, since even a model with poor performances can have a coverage of 100%: If its range is 1, it means that this model predicts that the occupancy probability is between 0 and 1.

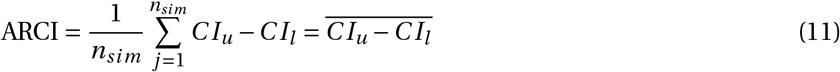

### 2.3 Empirical comparison with lynx detection data

We applied the five occupancy models to empirical data from Gimenez et al. (2022). We analyzed data from 11 camera-trap locations in Ain county, France, monitored from February 2017 to May 2019. Monitoring durations per site were heterogeneous, ranging from 148 to 801 days (Fig. S8), totaling 5396 camera-trap days across all 11 sites. We here focus on lynx (*Lynx lynx*) occupancy. Lynx were detected in 9 sites out of 11, with 203 detections in total, ranging from 1 to 59 detections per site (Fig. S8). For the two discrete models, BP and COP, we discretised the data similarly to the simulation study: into monthly, weekly, and daily intervals (Fig. S9). We discarded data from incomplete sessions, during which a site was monitored only partially and not throughout the entire session.

We estimated parameters following the method described in Section 2.2.2, consistent with the simulation study approach, with the exception of the optimisation algorithm starting points. We used intuitive starting points, such as the ratio of sites with at least one detection for *ψ*. Starting points for the models’ different detection parameters are detailled in our code. We did not include covariates in the analysis. For each parameter of each model, we retrieved its point estimate and derived its 95% and 50% confidence intervals from the Hessian matrix.

## 3 Results

### 3.1 Simulation study

For easily detectable species (detection scenarios d, e, f, and g), all models retrieve well the simulated occupancy probability. Bias ranges from *−*0.0094 to 0.0025 (Fig. 3) and RMSE are less than 0.060 (Fig. S2). With those detection parameters, the Kruskal-Wallis tests indicate that there are no statistically significant differences in the distribution of 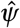 between models, except with simulation parameters (e) and *ψ =* 0.1, (e) and *ψ =* 0.9 and (f) and *ψ =* 0.9 (Table S3). The Wilcoxon tests indicate that there is no difference in medians with (e) and *ψ =* 0.1 (Fig. S3). With (e) and *ψ =* 0.9 and (f) and *ψ =* 0.9, only the BP model with daily sessions differs from the others, with a slight underestimation of *ψ* (Fig. 3).

**Figure 3:**
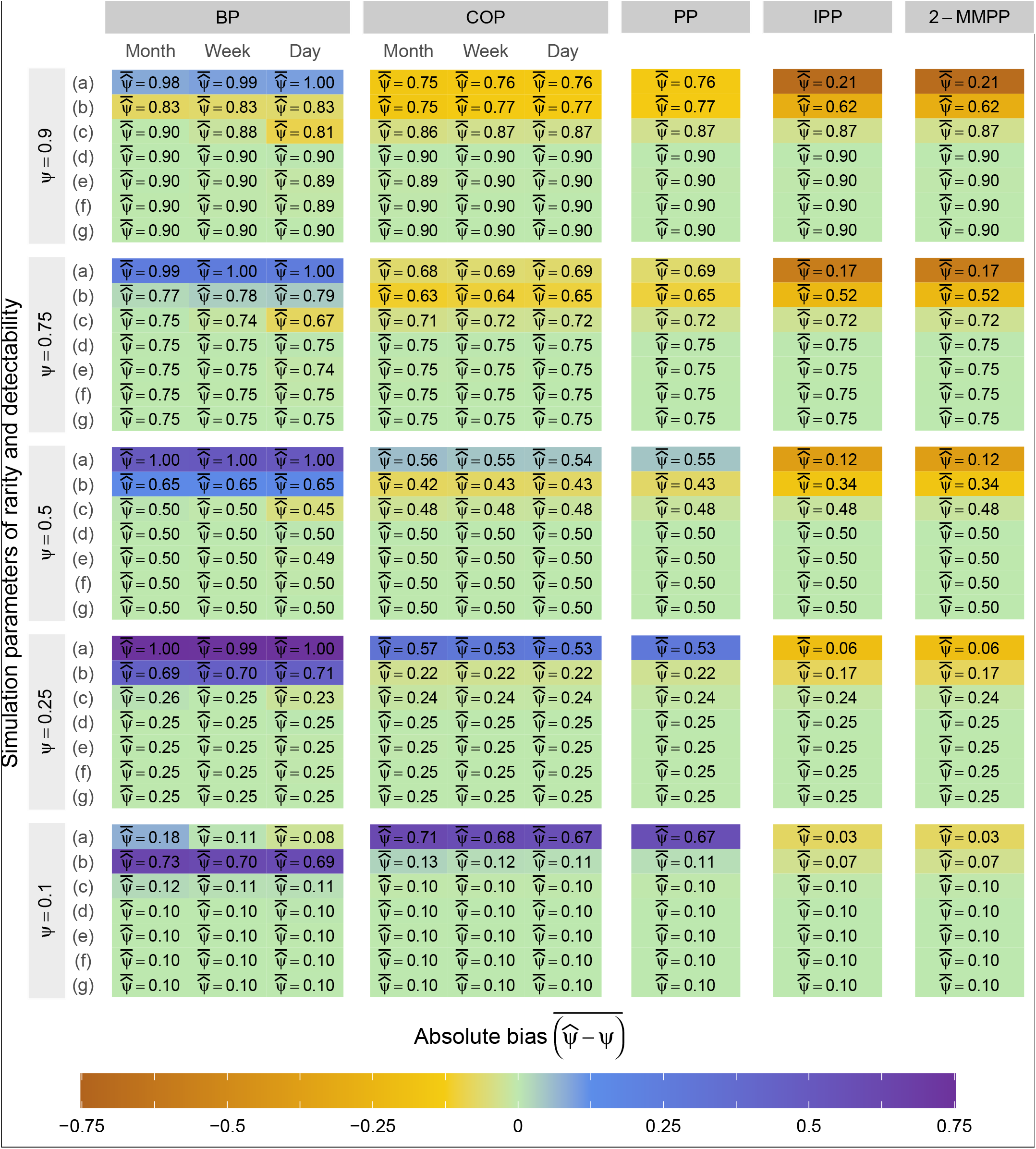
Absolute bias of the occupancy probability point estimate. Depending on *ψ* the simulated occupancy probability and detection scenarios as described in Table 2. The average value of the occupancy probability point estimate 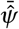 is inside each cell. For two scenarios characterised by low occupancy and detection probabilities, certain repetitions failed to yield any data. With no detection within any of the sites, it was impossible to infer parameters. With detection parameters (a) and *ψ =* 0.25, 494 simulations were used to estimate the models’ ability to retrieve the simulation parameters. With detection parameters (a) and *ψ =* 0.1, only 423 simulations were used.

For elusive species (detection scenario c), the BP model’s ability to retrieve the simulated occupancy probability is slightly inferior to other models, with a RMSE ranging from 0.057 to 0.121 while the RMSE of other models are still less than 0.060. (Fig. S2). The Wilcoxon tests (Fig. S3) indicate differences between BP and the other models, and this difference depends on the discretised session duration. The distribution of 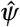 with BP is wider than for the other models with the same simulated data (Fig. S1).

For highly elusive species (detection scenarios a and b), all five models reach their limits. BP, COP and PP tend to overestimate *ψ*, whereas IPP and 2-MMPP tend to underestimate *ψ* (Fig. 3). BP tend to estimate *ψ* at 0 or most often at 1 (Fig. S1). COP and PP point estimates of *ψ* have similar distributions, both are widely spread (Fig. S1). IPP and 2-MMPP tend to underestimate *ψ*, with a tighter distribution for its point estimate, which often does not include the simulated value of *ψ* (Fig. S1).

It was not always possible to calculate the confidence interval (CI) of the occupancy probability estimate, when the Hessian matrix was not invertible. This occurred in two main cases in our study: when there were not many sessions with detections in the BP model, or when *λ*_1_ was estimated to zero in the 2-MMPP model. As a result, the 2-MMPP CIs are not interpretable with detection scenarios other than (d), where data were simulated as an IPP.

For easily detectable species (detection scenarios e, f and g), all models have similar coverages (Fig. S4) and occupancy probability CI ranges (Fig. S5). As detectability decreases, the CIs widens for BP, COP and PP, although this is more marked and quicker for BP than for COP and PP (Fig. S5). The IPP CIs do not widen, but the coverage drops (Fig. S4).

### 3.2 Application to lynx occupancy

Lynx occupancy estimates are similar across all five model (Fig. 4). The COP model, regardless of the session duration, and the three continuous models (PP, IPP, and 2-MMPP), converge to an estimated occupancy probability of 0.82, with identical confidence intervals (95% CI: 0.49 to 0.95, 50% CI: 0.73 to 0.88) (Fig. 4, Table S4). Although the BP model produced a slightly different point estimate, ranging between 0.75 and 0.80 depending on the session duration, it remains in close proximity to the estimates of the other four models (Fig. 4, Table S4).

**Figure 4:**
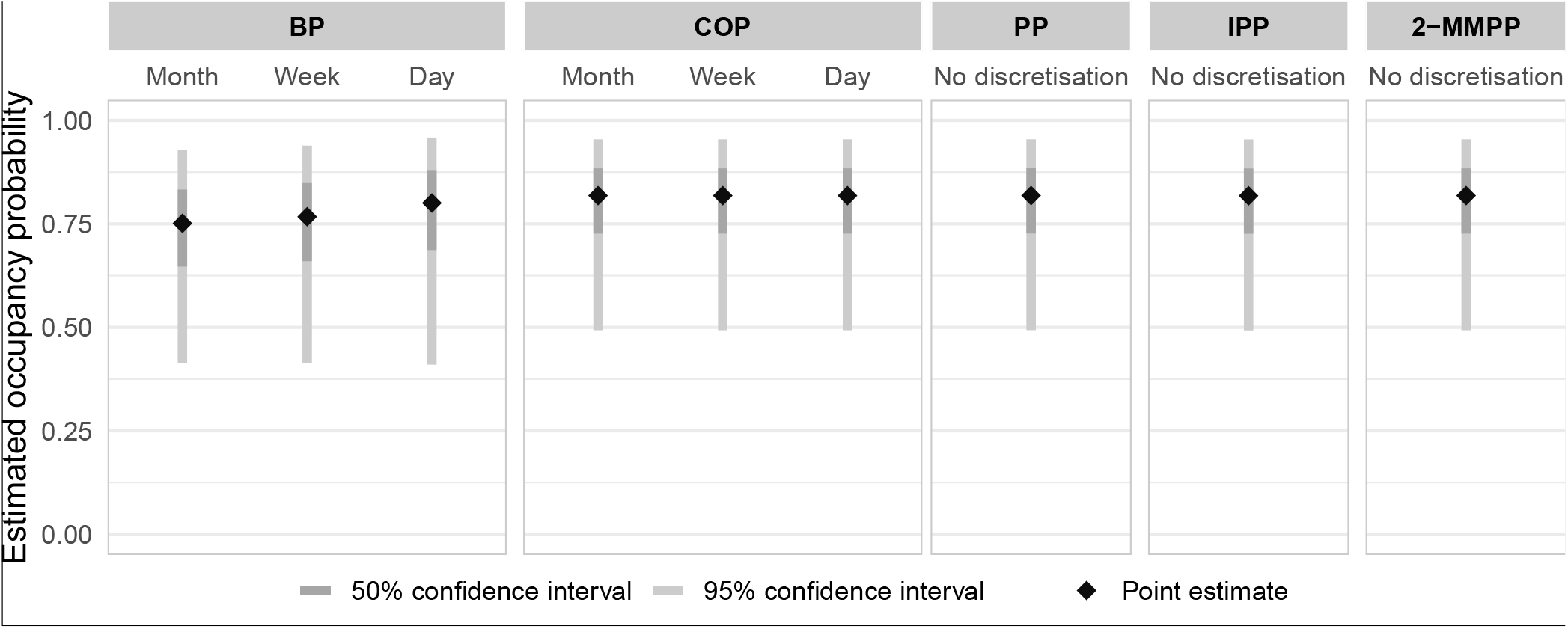
Occupancy probability estimation with the lynx data set from Gimenez et al. (2022) in the Ain County, France.

Regarding detectability, the BP model with daily sessions estimates the probability of detecting at least one lynx in a day at *p*_1_ *=* 0.008 (95% CI: 0.006 to 0.012, Table S4). Models assuming homogeneous detection rates (COP, PP) estimate 0.045 detection events per day (95% CI: 0.039 to 0.052, Table S4), resulting in an expected average of 𝔼 [*N*_100_] *=* 0.045 *×* 100 *=* 4.5 detection events in 100 days. In comparison to detectability in our simulation scenarios, lynx elusiveness likely falls between detection scenario (b) (*p*_1_ *=* 0.01) and scenario (c) (𝔼 [*N*_100_] *=* 6.25) (Table 2).

The 2-MMPP estimates a low detection rate in state 1, close to 0 (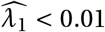, 95% CI: 0.001 to 0.06, Table S4), suggesting that lynx detection events can be modeled by the the IPP model. The IPP model estimates that only 0.31% of the time is spent in the state with detections, state 2, (Equation 4, with 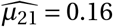 and 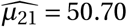, Table S4). The detection rate is estimated at 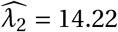 detection events per day in this state (95% CI: 9.24 to 21.89, Table S4).

## 4 Discussion

The focal ecological parameter of interest is the occupancy probability *ψ*, which is represented similarly in all the five models compared. However, the precision of the occupancy estimation is impacted by the quality of the estimation for the detection process (Kellner & Swihart, 2014; Kéry & Schmidt, 2008). In this study, we focused on cases in which data is collected continuously, for example with sensors or opportunistic data. We aimed to evaluate whether modelling the detection process in continuous-time could enhance the precision of the estimated probability of occupancy.

In line with the concept of operating models, commonly used for assessing management strategies (Butterworth, 1999; Punt et al., 2016), we simulated data under models that aim to get more in line with our expectations regarding the detection process when observing fauna through sensors. Specifically, we considered special cases of 2-MMPP, consisting of six scenarios with detections simulated under an IPP framework (scenarios a, b, c, e, f, g) and one scenario simulated under a PP framework (scenario d). We intentionally simulated occupancy very simply, not incorporating covariates or variations in abundance, aiming to focus solely on the detection process for sensor-collected data. Subsequently, we aimed to recover the simulation parameters, focusing on occupancy probability, using these complex models, as well as simpler models well-known and widely used by ecologists. By simplifying the information and the detection process, we asked the question of whether these models are sufficient to estimate the ecological parameter of interest in a situation that we expect to be close to reality.

We expected that continuous models would outperform discrete models in accurately retrieving the simulated occupancy probability, since detection data simulation aligned with the framework of the continuous models of our comparison. Moreover, data aggregation by discretisation leads to a loss of information. We expected this loss to result in less accurate estimates of detection parameters, consequently introducing bias in the occupancy estimate, given the relationship between occupancy and detection parameters. However, in the majority of cases where detectability was sufficiently high (with a minimum expectation of 25 detections in occupied sites throughout the entire deployment), all models produced equivalent results, all were able to retrieve the occupancy probability well, with little bias and error.

For models requiring discrete data, we expected that different discretisations would impact the models outputs (Schofield et al., 2018), but in most simulated scenarios, that was not the case. Our results indicate that estimation of *ψ* with BP is more impacted by the session duration’s choice than with COP. Since COP is mathematically equivalent to PP (Zhang & Bonner, 2020), minor variations in the occupancy estimates between session lengths for COP are likely due to data discarding. Our comparison framework could be reused to further test the impact of discretisation, by choosing more diverse session durations that reuse exactly the same data - rather than dropping data of incomplete sessions as we did.

The BP model, as noted by MacKenzie et al. (2002), tends to produce estimates of *ψ* close to one for rare and elusive species. Our findings align with this observation, suggesting however that elusiveness has a more pronounced impact on this limit than rarity.

The COP model was adapted from the model proposed by Emmet et al. (2021). Their model differs from the one presented here mainly because they considered site use. However, they compared their counting model with its detection/non-detection equivalent from Bled et al. (2013), much like we compared COP with BP. Their model estimated occupancy probability with either equivalent or smaller bias compared to the equivalent detection/non-detection model, which aligns with our results.

In a simulation study, Guillera-Arroita et al. (2011) evaluated BP and PP using data generated within a PP framework. They reported that both models provided reasonably unbiased estimates of occupancy, except for rare and elusive species. In these cases, BP exhibited greater bias and variance, particularly with larger discretisation intervals and fewer sessions, which matches our results. They also compared PP and 2-MMPP using clustered detection data generated within an IPP framework. They noted negative bias in the occupancy estimates with the PP model, which was not observed in our results. In our study, both models performed similarly for easily detectable species. However, for elusive species, the 2-MMPP and IPP models exhibited more pronounced negative bias than the PP and COP models.

All five models consistently estimated lynx occupancy in our empirical example, with COP, PP, IPP, and 2-MMPP estimating occupancy probability at 0.82, while BP provided slightly lower but still very close estimates. Lynx detectability falls between simulation scenarios (b) and (c), where models began to approach their limits due to elusiveness in our simulations. As different models produced significantly varied occupancy estimates when they reached their limits in simulations, the consistent occupancy estimates across models suggest that they have not yet reached their limits, indicating reliable occupancy estimates.

To better define the limitations of these models, we could perform additional comparisons using simulation scenarios with various detection parameters. Given the impossibility of exhaustively covering all potential scenarios, we encourage modelers to compare models when encountering borderline cases of occupancy models applicability, such as potentially insufficient monitoring time in view of the species elusiveness. Our code is available to use as a base to conduct simulations with parameters adapted to a specific study context and compare models to choose the best model. Alternatively, researchers and practitioners can analyse their data with multiple models to ensure the consistency of results across different modelling approaches, as we did with the lynx data set.

### 4.1 Choosing the appropriate model

#### 4.1.1 Occupancy modelling for easily detectable species

When the species is easily detectable and thus enough observation data have been obtained, all models accurately estimate the occupation probability. Under these conditions, if the sole aim of a study is to accurately estimate occupancy, selecting any of these models essentially amounts to choosing the right one. Therefore, the choice can be guided by other considerations, to find the right balance between performance and execution costs.

##### Learning and implementation costs

Continuous-time models may be unfamiliar to ecologists, potentially requiring a steep learning curve to become proficient with these seemingly complex models. For models that are not readily available, the implementation costs can be substantial for a study. The rise of sensor-based monitoring and the growing interest have led to efforts to make time-to-detection occupancy models more accessible for ecologists, such as through R packages like `unmarked` (Kellner et al., 2023). The costs shifts from fully implementing a model to using existing functions, a faster alternative. We are currently working on adding the COP, PP, IPP and 2-MMPP models to `unmarked`.

##### Study objectives

If the primary goal is to estimate the occupancy of the target species, any of the models can be employed effectively. Simple models, such as BP, COP or PP, require the estimation of only two parameters: one for occupancy and one for detection. Choosing such a model can enhance interpretability and provide a greater statistical power than models with more parameters. This is especially advantageous when incorporating several spatial and temporal covariates into the analysis. Conversely, if the aim is to conduct a detailed analysis of the target species detection timeline to better understand a species behaviour, analysing continuous-time data is preferable, as it data holds valuable information that is increasingly lost as data are more and more aggregated over time. Models that accommodate the detection process in multiple states could be particularly adapted to unravel animal behaviours, such as temporal activity patterns. The hidden state could be reconstructed to further enhance the interpretability of these models’ parameters.

##### Temporal auto-correlation

Unlike sampling occasions, consecutive discretised sessions are not temporally independent (Bailey et al., 2014), and there may be significant temporal auto-correlation (Neilson et al., 2018). Therefore, discretised session data does not meet the discrete-time model assumption of independence. However, the PP model has the exact same drawback when considering a constant detection rate, since the number of events on two disjoints time intervals are independent. In this study, we did not thoroughly examine the influence of time dependence on occupancy estimates. However, two-state models, introduced for clustered observation data (Guillera-Arroita et al., 2011), incorporate some time dependence through two homogeneous detection rates that differ conditional on state. Future studies could explore the impact of time dependence on occupancy estimates and consider various approaches to account for non-constant detection rates. These may include Cox processes, where the detection rate is a random variable (Cox, 1955), time-dependent regression splines (Distiller et al., 2020), or Hawkes processes, a form of self-exciting point process where the detection rate increases temporarily after a detection event (Hawkes, 1971; Rushing, 2023).

##### Calculation time

All models were fairly fast to fit, so calculation time should probably not be the main reason for choosing a model for most studies. We have not robustly evaluated the optimisation time for each model, as we used different computers with varying characteristics. However, the two-state models seemed significantly longer to fit than the other models. BP, COP and PP all took less than 6 seconds to fit, even in the simulation scenario with most detections, in which there was 200 detections on average in occupied sites. IPP and 2-MMPP often took more than a minute, up to 28 minutes.

##### Detection rate

A detection probability per discretised session, as in the BP model, is relevant only at the discretisation scale. This is not the case with a detection rate, as used in the discrete-time COP model or in continuous-time models. We argue that using a detection rate instead of a detection probability would enhance the comparability among studies. This could simplify the process of experimental design, especially concerning observation duration, by using the insights from existing literature on the target species. Another advantage of using a detection rate, as opposed to a probability, is the flexibility to accommodate sessions of different durations. For instance, in our lynx example, monthly sessions varied in length (28, 30, or 31 days). We specified this in the COP model to estimate the number of detection per day, an explicit time unit. We could not do so in the BP model, in which the detection probability is derived per session.

#### 4.1.2 Occupancy modelling for highly elusive species

When the species is highly elusive, the five models we considered provided inaccurate estimates of its presence probability, exhibiting high bias, error, and a low precision or coverage. The BP model’s limits became apparent at lower species elusiveness compared to the other models. This could be because valuable information gets lost when simplifying the data into detection and no detection. The 2-MMPP and IPP models showed larger errors in estimating *ψ* compared to the COP and PP models. This might be due to the higher number of parameters in the 2-MMPP and IPP models (5 and 4, respectively, versus 2 for COP and PP), which would require more data to fit them correctly. COP and PP models appear to strike a good balance between simplification and realism. One is discrete, while the other is continuous, but both perform similarly, which is consistent with the demonstration of Zhang and Bonner (2020) that a Poisson process in continuous time is equivalent to a classical model with discretisation where the detection process is not a Bernoulli trial but a Poisson distribution draw.

However, if the species’ high elusiveness resulted in the collection of insufficient observation data, the best course of action probably is to collect more data by extending the monitoring period (Kays et al., 2020). In cases where it is expected that the species will be challenging to detect, conducting simulations and comparing different models with expected detection and occupancy parameters could assist in fine-tuning the study design and model choice.

If obtaining more data is not feasible, it might be best to refrain from running an occupancy model, or at least approach the results with caution, regardless of the chosen model. In this case, we recommend fitting different models, particularly when using the two-state models. For these models, our findings indicate that with highly elusive species, the confidence interval of the estimated *ψ* can be narrow but substantially different from the actual *ψ*. This can potentially lead to a misleading perception of model reliability.

### 4.2 Implications for continuous monitoring frameworks

The advanced processors available today offer great computing power, enabling the fast development of artificial intelligence (AI). Recognising species automatically is becoming more common, on camera-trap images (Le Borgne & Bouget, 2023), ARUs recordings (Potamitis et al., 2014), or even with sensors networks (Wägele et al., 2022). AI combined with sensors offers the potential to fully automate the analysis workflow (Gimenez et al., 2022; Lahoz-Monfort & Magrath, 2021). Overall, sensors and AI have led to a paradigm shift in the conditions and capabilities of biodiversity monitoring (Besson et al., 2022; Tuia et al., 2022; Zwerts et al., 2021). With our comparison, we found that modelling occupancy with a continuous-time detection sub-model is not necessary to estimate occupancy accurately. Therefore, in operational conditions, the necessary trade-off between accuracy and ease of implementation may turn in favour of discrete-time models, with easily available data for temporal covariates.

Focusing primarily on the temporal aspect of the detection process modelling, we explored a simple version of occupancy models: single-species, static, with no positive and no abundance-induced heterogeneity. There are numerous avenues for further investigation. With the emergence of collaborative platforms aggregating and providing sensor data across large spatiotemporal scales (e.g. Oliver et al., 2023), developing dynamic occupancy models with continuous-time occupancy sub-models has the potential to enhance our understanding of extinction-colonisation processes, to get the most out of continuous-time data. Additionally, we did not consider abundance as a factor influencing detection heterogeneity, although it is a particularly intuitive consideration for species with highly variable abundance across sites (Royle & Nichols, 2003). Further research into these models, in conjunction with continuous data, is called for: In N-mixture models, time-to-first-detection and time-to-each-detection could offer improved estimates compared to binary detection/non-detection and count data, respectively (Haines et al., 2023; Priyadarshani et al., 2024). Furthermore, our study did not account for false-positives, and sensor data (e.g., images from camera traps) can be prone to incorrect species identification. Potential solution, such as using AI confidence scores (Rhinehart et al., 2022), could be explored further with continuous-time data to consider the complete workflow, from sensor data to inference.

Our results do not only concern sensor data, but all continuously collected data. Opportunistic data, collected at non-defined and irregular intervals, can be considered as continuously collected and thus pose some of the same challenges as sensor data (Altwegg & Nichols, 2019; Hsing, 2019). Some studies analyse opportunistic data using with discrete-time models, discretising data into long sessions (*e*.*g*., by year, as in van Strien et al., 2013), while other are developing new continuous-time frameworks adapted to this particular data (*e*.*g*. Choquet et al., 2017, using continuous-time capture-recapture for individualised opportunistic data). Our comparison suggests that for future studies aiming to estimate occupancy with unmarked opportunistic data, both discrete- and continuous-time occupancy models could produce accurate occupancy estimates, provided that challenges associated with opportunistic data, such as highly variable observation effort, are effectively addressed.

## Acknowledgements

LP benefits from a French government CIFRE grant for PhD students. This work is part of the PSI-BIOM project granted by the French PIA 3 under grant number 2182D0406-A.

We thank the Federations of Hunters from the Jura and Ain counties for sharing the lynx camera trap data. Data collection was carried out through the Lynx Predator Prey Program, which was funded by Auvergne-Rhone-Alpes Region, Ain and Jura departmental Councils, the French National Federation of Hunters, French Environmental Ministry based in Auvergne-Rhone-Alpes and Bourgogne-Franche-Comte Region and the French Office for Biodiversity.

## Author Contributions

- LP: Formal analysis, Investigation, Methodology, Visualisation, Writing – original draft.
- SM, OG, MPE: Validation, Writing – review and editing.
- All authors: Conceptualisation.

## Code availability

Code is available at: https://oikolab.terroiko.fr:10001/publications/occupancy-modelling-comparison-discrete-continuous

