## Supplementary information for "Analysing biodiversity observation data collected in continuous time: Should we use discrete- or continuous-time occupancy models?"

#### Contents

Formulas, parameters and associated publications

Counting Occurrences Process Likelihood

Mean and variance of the number of detections for a 2-MMPP

Estimation with Nelder-Mead: Supplementary figures

Estimation with BFGS

Application to lynx occupancy

### Appendix 1 Formulas, parameters and associated publications

Table S1: **Formulas, parameters and associated publications**. All mathematical notation are summarised in Table 1 of the manuscript.

(a) Models overview

| Model | Reference | Time | Data | Parameters |
|---|---|---|---|---|
| **BP** | MacKenzie et al. (2002) | Discrete | $Y_{is} \in \{0;1\}$ | $\psi$, $p$ |
| **COP** | Emmet et al. (2021), Appendix 2 | Discrete | $N_{is} \in \mathbb{N}$ | $\psi$, $\lambda$ |
| **PP** | Guillera-Arroita et al. (2011) | Continuous | $t_{i1}, \ldots, t_{iN_i}$ | $\psi$, $\lambda$ |
| **2-MMPP** | Guillera-Arroita et al. (2011) | Continuous | $t_{i1}, \ldots, t_{iN_i}$ | $\psi, \lambda_1, \lambda_2, \mu_{12}, \mu_{21}$ |
| **IPP** | Guillera-Arroita et al. (2011) | Continuous | $t_{i1}, \ldots, t_{iN_i}$ | $\psi, \lambda_2, \mu_{12}, \mu_{21}$ |

(b) Formulas, with # the equation number in the manuscript

| | # | Formula | Reference |
|---|---|---|---|
| **All models**<br>Occupancy sub-model | (1) | $Z_i \sim \text{Bernoulli}(\psi)$ | MacKenzie et al. (2002) |
| **BP**<br>Detection sub-model | (2) | $Y_{is} \vert Z_i = 1 \sim \text{Bernoulli}(p)$<br>$Y_{is} \vert Z_i = 0 \sim 0$ | MacKenzie et al. (2002) |
| **COP**<br>Detection sub-model | (3) | $N_{is} \vert Z_i = 1 \sim \text{Poisson}((\lambda T_s)$<br>$N_{is} \vert Z_i = 0 \sim 0$ | Emmet et al. (2021), Appendix 2 |
| **2-MMPP**<br>Steady state vector | (4) | $\Pi = \begin{pmatrix} \pi_1 & \pi_2 \end{pmatrix} = \begin{pmatrix} \frac{\mu_{21}}{\mu_{12}+\mu_{21}} & \frac{\mu_{12}}{\mu_{12}+\mu_{21}} \end{pmatrix}$ | Fischer and Meier-Hellstern (1993) |
| **2-MMPP**<br>Number of detections | (5)<br>(6) | $\mathbb{E}[N_{it}] = (\lambda_1\pi_1 + \lambda_2\pi_2)\,t$<br>$\mathbb{V}[N_{it}] = \left(\lambda_1\pi_1 + \lambda_2\pi_2 + \frac{2({\lambda_1}^2+{\lambda_2}^2)}{{\mu_{12}}^2{\mu_{21}}^2\left(\frac{1}{\mu_{12}}+\frac{1}{\mu_{21}}\right)^3}\right) t$ | Bhat (1992), Appendix 3 |
| **2-MMPP**<br>Detection probability | (7) | $p_t = 1 - \Pi exp\left[\left(\begin{pmatrix} -\mu_{12} & \mu_{12} \\ \mu_{21} & -\mu_{21} \end{pmatrix} - \begin{pmatrix} \lambda_1 & 0 \\ 0 & \lambda_2 \end{pmatrix}\right) t\right] \begin{pmatrix} 1 \\ 1 \end{pmatrix}$ | Guillera-Arroita et al. (2011) |
| **Evaluation metrics**<br>$\psi$ point estimate ($\hat{\psi}$) | (8)<br>(9) | $\text{RMSE} = \sqrt{\frac{1}{n_{sim}} \sum_{j=1}^{n_{sim}} \left(\hat{\psi}_j - \psi\right)^2} = \sqrt{\overline{\left(\hat{\psi} - \psi\right)^2}}$<br>$\text{AB} = \frac{1}{n_{sim}} \sum_{j=1}^{n_{sim}} (\hat{\psi}_j - \psi) = \overline{(\hat{\psi} - \psi)}$ | See for example<br>Liemohn et al. (2021) |
| **Evaluation metrics**<br>$\psi$ confidence interval<br>$[CI_l, CI_u]$ | (8)<br>(9) | $\text{Coverage} = \frac{1}{n_{sim}} \sum_{j=1}^{n_{sim}} I(CI_l \leq \psi \leq CI_u) = \overline{I(CI_l \leq \psi \leq CI_u)}$<br>$\text{ARCI} = \frac{1}{n_{sim}} \sum_{j=1}^{n_{sim}} CI_u - CI_l = \overline{CI_u - CI_l}$ | See for example<br>Liemohn et al. (2021)<br>Schall (2012) |

### Appendix 2 Counting Occurrences Process Likelihood

The COP model is based on the one proposed by (Emmet et al., 2021). Hereafter, we use the notation defined in Table S2.

Table S2: **Mathematical notation for the COP model**.

| | |
|---|---|
| $I$ | Number of sites |
| $S$ | Number of sessions |
| $\psi_i$ | Occupancy probability in site $i$ |
| $Z_i$ | Occupancy state of site $i$ (present = 1, absent = 0) |
| $\lambda_{is}$ | Detection rate in site $i$ during session $s$ |
| $T_{is}$ | Duration of session $s$ in site $i$ |
| $N_{is}$ | Number of detection events in site $i$ during session $s$ |
| $N_i$ | Total number of detection events in site $i$ |

The COP model can be described with Equation Appendix 2.1, with $Z_i$ the occupancy state of site $i$; $\psi_i$ the occupancy probability of site $i$; $N_{is}$ the number of detections in site $i$ during session $s$; $\lambda_{is}$ the detection rate in site $i$ during session $s$; and $T_{is}$ the duration of session $s$ in site $i$.

$$\begin{aligned} Z_i &\sim Bernoulli(\psi_i) \\ N_{is} &\sim Poisson(\lambda_{is} T_{is}) \end{aligned} \qquad \text{(Appendix 2.1)}$$

##### Probability of having $N_i = n_i$ detection events in a site

We consider $N_{i1}, N_{i2}, \ldots, N_{iS}$ to be independent Poisson random variables. The sum of random Poisson variables is another Poisson, with the sum of their rates as its rate. Therefore, instead of calculating the probability of having the $N_{is} = n_{is}$ for each session $s$ in site $i$, we can set $n_i = \sum_{s=1}^{S} n_{is}$ and calculate the probability of having $N_i = n_i$, the total number of detections in site $i$, with $N_i \sim Poisson\left(\sum_{s=1}^{S}(\lambda_{is} T_{is})\right)$.

The probability of having $n_i$ detection events can be splitted into two cases, depending on wether if the site is occupied ($z_i = 1$) or not ($z_i = 0$):

$$\mathbb{P}_{\psi_i,\lambda_{is}}(N_i = n_i) = \mathbb{P}_{\psi_i,\lambda_{is}}(N_i = n_i, z_i = 1) + \mathbb{P}_{\psi_i,\lambda_{is}}(N_i = n_i, z_i = 0)$$

For each site, will calculate this probability in two cases, depending on if there were detections ($n_i > 0$) or not ($n_i = 0$) in site $i$.

###### In a site with at least one detection event

$$\mathbb{P}_{\psi_i,\lambda_{is}}(N_i = n_i, n_i > 0) = \mathbb{P}_{\psi_i,\lambda_{is}}(N_i = n_i, n_i > 0, z_i = 1) + \mathbb{P}_{\psi_i,\lambda_{is}}(N_i = n_i, n_i > 0, z_i = 0)$$

If there is at least one detection in site $i$ ($n_i > 0$), the site $i$ is considered occupied. Another way to see it is that if the species does not occupy the site $i$ ($z_i = 0$), it is not possible to detect it, hence $\mathbb{P}_{\psi_i,\lambda_{is}}(N_{is} = n_{is}, n_i > 0, z_i = 0) = 0$. Therefore, we have:

$$\begin{aligned}\mathbb{P}_{\psi_i,\lambda_{is}}(N_i = n_i, n_i > 0) &= \mathbb{P}_{\psi_i,\lambda_{is}}(N_i = n_i, n_i > 0, z_i = 1)\\ &= \mathbb{P}_{\psi_i,\lambda_{is}}(z_i = 1)\mathbb{P}_{\psi,\lambda}(N_i = n_i, n_i > 0)\\ &= \psi_i \mathbb{P}_{\psi,\lambda}(N_i = n_i, n_i > 0)\end{aligned}$$

Because $N_i \sim Poisson\left(\sum_{s=1}^{S}(\lambda_{is}T_{is})\right)$, we therefore have:

$$\mathbb{P}_{\psi_i,\lambda_{is}}(N_i = n_i, n_i > 0) = \psi_i \frac{\left(\sum_{s=1}^{S}(\lambda_{is}T_{is})\right)^{n_i}}{n_i!} e^{-\sum_{s=1}^{S}(\lambda_{is}T_{is})} \tag{Appendix 2.2}$$

**Without covariates, and with constant session durations, we can simplify this equation.**

Without site covariates, all sites have the same occupancy probability that we will write $\psi$ ($\psi = \psi_i,\ i = 1,...,I$). Without temporal covariates, all sites and all sessions have the same detection rate that we will write $\lambda$ ($\lambda = \lambda_{is},\ i = 1,...,I,\ s = 1,...,S$).

We also set $T_s$ the duration of a session, with all sessions in all sites constant ($T_s = T_{is},\ i = 1,...,I,\ s = 1,...,S$), and $S$ the number of sessions that were carried out in site $i$. We therefore have $ST_s = \sum_{s=1}^{S} T_{is}$.

With those simplifications, we have:

$$\mathbb{P}_{\psi,\lambda}(N_i = n_i, n_i > 0) = \psi \frac{(\lambda ST_s)^{n_i}}{n_i!} e^{-\lambda ST_s} \tag{Appendix 2.3}$$

**In a site with no detection**

Having no detection in a site can happen in two situations: (1) the site is occupied but the species was not detected, or (2) the site is not occupied.

$$\begin{aligned}\mathbb{P}_{\psi_i,\lambda_{is}}(N_i = 0) &= \mathbb{P}_{\psi_i,\lambda_{is}}(N_i = 0, z_i = 1) + \mathbb{P}_{\psi_i,\lambda_{is}}(N_i = 0, z_i = 0)\\ &= \mathbb{P}_{\psi_i,\lambda_{is}}(z_i = 1) \times \mathbb{P}_{\psi_i,\lambda_{is}}(N_i = 0 \mid z_i = 1) + \mathbb{P}_{\psi_i,\lambda_{is}}(z_i = 0) \times \mathbb{P}_{\psi_i,\lambda_{is}}(N_i = 0 \mid z_i = 0)\end{aligned}$$

We have $\mathbb{P}_{\psi_i,\lambda_{is}}(N_i = 0 \mid z_i = 0) = 1$, because if the site is not occupied, we can not detect the species, so the probability of having no detection events is 1. Moreover, $N_i \sim Poisson\left(\sum_{s=1}^{S}(\lambda_{is}T_{is})\right)$, so $\mathbb{P}_{\psi_i,\lambda_{is}}(N_i = 0 \mid z_i = 1) = \frac{\left(\sum_{s=1}^{S}(\lambda_{is}T_{is})\right)^{n_i}}{n_i!} e^{-\sum_{s=1}^{S}(\lambda_{is}T_{is})}$. But because $n_i = 0$, this term can be simplified to $e^{-\sum_{s=1}^{S}(\lambda_{is}T_{is})}$. Therefore, we have:

$$\mathbb{P}_{\psi_i,\lambda_{is}}(N_i = 0) = \psi_i e^{-\sum_{s=1}^{S}(\lambda_{is}T_{is})} + (1 - \psi_i) \tag{Appendix 2.4}$$

**Without covariates, and with constant session durations, we can simplify this equation**, like we did in the precedent section:

$$\mathbb{P}_{\psi_i,\lambda_{is}}(N_i = 0) = \psi_i e^{-\lambda ST_s} + (1 - \psi_i) \tag{Appendix 2.5}$$

##### Likelihood for all sites

The likelihood is the product of the probabilities in Equation Appendix 2.2 and Equation Appendix 2.4 for all sites.

$$
\begin{aligned}
L(\psi_i,\lambda_{is}) &= \prod_{i=1}^{I} \mathbb{P}_{\psi_i,\lambda_{is}}(N_i = n_i) \\
&= \prod_{i,n_i>0} \left(\mathbb{P}_{\psi_i,\lambda_{is}}(N_i = n_i, n_i > 0)\right) \times \prod_{i,n_i=0} \left(\mathbb{P}_{\psi_i,\lambda_{is}}(N_i = 0)\right) \\
&= \prod_{i,n_i>0} \left(\psi_i \frac{\left(\sum_{s=1}^{S}(\lambda_{is}T_{is})\right)^{n_i}}{n_i!} e^{-\sum_{s=1}^{S}(\lambda_{is}T_{is})}\right) \times \prod_{i,n_i=0} \left(\psi_i e^{-\sum_{s=1}^{S}(\lambda_{is}T_{is})} + (1-\psi_i)\right)
\end{aligned}
\tag{Appendix 2.6}
$$

**Without covariates, and with constant session durations, we can simplify the likelihood**, using Equation Appendix 2.3 and Equation Appendix 2.5. We note $S_+$ the number of sites where there was at least a detection, $n$ the total number of detections, $S_{tot}$ the total number of sessions of length $T_s$, the likelihood for this model is:

$$
\begin{aligned}
L(\psi,\lambda) &= \prod_{i=1}^{I} \mathbb{P}_{\psi,\lambda}(N_i = n_i) \\
&= \prod_{i,n_i>0} \left(\mathbb{P}_{\psi,\lambda}(N_i = n_i, n_i > 0)\right) \times \prod_{i,n_i=0} \left(\mathbb{P}_{\psi,\lambda}(N_i = 0)\right) \\
&= \prod_{i,n_i>0} \left(\psi \frac{(S\lambda T_s)^{n_i}}{n_i!} e^{-S\lambda T_s}\right) \times \prod_{i,n_i=0} \left((1-\psi) + \psi \times e^{-S\lambda T_s}\right) \\
&= \left(\psi^{S_+} e^{-S_+ S_{tot}\lambda T_s} (S_{tot}\lambda T_s)^n \prod_{i,n_i>0}\left[\frac{1}{n_i!}\right]\right) \times \left((1-\psi) + \psi e^{-S_{tot}\lambda T_s}\right)^{S-S_+}
\end{aligned}
\tag{Appendix 2.7}
$$

### Appendix 3 Mean and variance of the number of detections for a 2-MMPP

With :

- $s_1$ the random variable of the time spent in state 1 before switching ($s_2$ in state 2).
  - $s_1 \sim \text{Exp}(\mu_{12})$ and $s_2 \sim \text{Exp}(\mu_{21})$
  - $\mathbb{E}[s_1] = 1/\mu_{12}$ and $\mathbb{E}[s_2] = 1/\mu_{21}$
  - $\mathbb{V}[s_1] = 1/\mu_{12}{}^2$ and $\mathbb{V}[s_2] = 1/\mu_{21}{}^2$
- $Z_{1t}$ the random variable of the time spent in state 1 during time $t$ ($Z_{2t}$ for state 2)
- $N_t^{S=1}$ the number of detections during the time-interval $t$ given that the system is in state 1 ($N_t^{S=2}$ when in state 2)

Using properties defined by Cox and Miller (1977), Equation 3 from Bhat (1992) gives the mean time spent in a state during a time period $t$:

$$\mathbb{E}[Z_{1t}] = \frac{\mathbb{E}[s_1]}{\mathbb{E}[s_1] + \mathbb{E}[s_2]} = \frac{\mu_{21}}{\mu_{12} + \mu_{21}} t + o(1) \approx \pi_1 t \tag{Appendix 3.1}$$

$$\mathbb{E}[Z_{2t}] = \frac{\mathbb{E}[s_2]}{\mathbb{E}[s_1] + \mathbb{E}[s_2]} = \frac{\mu_{12}}{\mu_{12} + \mu_{21}} t + o(1) \approx \pi_2 t \tag{Appendix 3.2}$$

Equation 4 (Bhat, 1992) gives the variance of time spent in a state during a time period $t$:

$$\begin{aligned}
\mathbb{V}[Z_{1t}] = \mathbb{V}[Z_{2t}] &= \frac{\mathbb{E}[s_1]^2 \mathbb{V}[s_2] + \mathbb{E}[s_2]^2 \mathbb{V}[s_1]}{(\mathbb{E}[s_1] + \mathbb{E}[s_2])^3} t + o(1) \\
&= \frac{\left(\frac{1}{\mu_{12}}\right)^2 \times \frac{1}{\mu_{21}{}^2} + \left(\frac{1}{\mu_{21}}\right)^2 \times \frac{1}{\mu_{12}{}^2}}{\left(\frac{1}{\mu_{12}} + \frac{1}{\mu_{21}}\right)^3} t + o(1) \\
&\approx \frac{2}{\mu_{12}{}^2 \mu_{21}{}^2 \left(\frac{1}{\mu_{12}} + \frac{1}{\mu_{21}}\right)^3} t
\end{aligned} \tag{Appendix 3.3}$$

Equation 5 from Bhat (1992) gives the expected number of events happening during a time period $t$, given that the system is in state 1 or 2 during $t$. Using Equation Appendix 3.1 and Equation Appendix 3.2, we have:

$$\begin{aligned}
\mathbb{E}[N_t^{S=1}] &= \lambda_1 \frac{\mathbb{E}[s_1]}{\mathbb{E}[s_1] + \mathbb{E}[s_2]} t + o(1) \\
&\approx \lambda_1 \pi_1 t
\end{aligned} \tag{Appendix 3.4}$$

$$\begin{aligned}
\mathbb{E}[N_t^{S=2}] &= \lambda_2 \frac{\mathbb{E}[s_2]}{\mathbb{E}[s_1] + \mathbb{E}[s_2]} t + o(1) \\
&\approx \lambda_2 \pi_2 t
\end{aligned} \tag{Appendix 3.5}$$

Equation 6 from Bhat (1992) gives the variance of the number of events happening during a time period $t$, given that the system is in state 1 or 2 during $t$. Using Equation Appendix 3.1, Equation Appendix 3.2 and Equation Appendix 3.3, we have:

$$\begin{aligned}\mathbb{V}[N_t^{S=1}] &= \mathbb{E}[Z_{1t}]\lambda_1 + \mathbb{V}[Z_{1t}]{\lambda_1}^2 \\ &\approx \left(\lambda_1\pi_1 + \frac{2}{{\mu_{12}}^2{\mu_{21}}^2\left(\frac{1}{\mu_{12}}+\frac{1}{\mu_{21}}\right)^3}{\lambda_1}^2\right)t \qquad \text{(Appendix 3.6)} \\ \mathbb{V}[N_t^{S=2}] &= \mathbb{E}[Z_{2t}]\lambda_2 + \mathbb{V}[Z_{2t}]{\lambda_2}^2 \\ &\approx \left(\lambda_2\pi_2 + \frac{2}{{\mu_{12}}^2{\mu_{21}}^2\left(\frac{1}{\mu_{12}}+\frac{1}{\mu_{21}}\right)^3}{\lambda_2}^2\right)t \qquad \text{(Appendix 3.7)}\end{aligned}$$

Since the number of detections in state 1 and in state 2 are independent, $N_t$ the random variable of the **total number of detections during a time period** $t$ is characterised by:

$$\begin{aligned}\mathbb{E}[N_t] &= \mathbb{E}[N_t^{S=1} + N_t^{S=2}] \\ &= \mathbb{E}[N_t^{S=1}] + \mathbb{E}[N_t^{S=2}] \\ &= (\lambda_1\pi_1 + \lambda_2\pi_2)\,t \qquad \text{(Appendix 3.8)} \\ \mathbb{V}[N_t] &= \mathbb{V}[N_t^{S=1} + N_t^{S=2}] \\ &= \mathbb{V}[N_t^{S=1}] + \mathbb{V}[N_t^{S=2}] \\ &= \left(\lambda_1\pi_1 + \lambda_2\pi_2 + \frac{2}{{\mu_{12}}^2{\mu_{21}}^2\left(\frac{1}{\mu_{12}}+\frac{1}{\mu_{21}}\right)^3}({\lambda_1}^2 + {\lambda_2}^2)\right)t \qquad \text{(Appendix 3.9)}\end{aligned}$$

#### Appendix 4 Estimation with Nelder-Mead: Supplementary figures

Table S3: **Kruskall-Wallis test results for simulation scenario of occupancy ($\psi$) and detection (as described in ??).** Presented with the Kruskal-Wallis rank sum statistic and the corresponding p-value. We compare nine groups (BP-month, BP-week, BP-day, COP-month, COP-week, COP-day, PP, IPP, and 2-MMPP) based on the distribution of the point estimate of the occupancy probability.

| | $\psi = 0.1$ | $\psi = 0.25$ | $\psi = 0.5$ | $\psi = 0.75$ | $\psi = 0.9$ |
|---|---|---|---|---|---|
| (a) | 2695.46<br>p < 2e-16 (***) | 2210.50<br>p < 2e-16 (***) | 2810.67<br>p < 2e-16 (***) | 2697.74<br>p < 2e-16 (***) | 2586.81<br>p < 2e-16 (***) |
| (b) | 884.45<br>p < 2e-16 (***) | 1127.39<br>p < 2e-16 (***) | 972.52<br>p < 2e-16 (***) | 1405.05<br>p < 2e-16 (***) | 1503.94<br>p < 2e-16 (***) |
| (c) | 53.88<br>p = 7.3e-09 (***) | 86.21<br>p = 2.7e-15 (***) | 169.74<br>p < 2e-16 (***) | 342.59<br>p < 2e-16 (***) | 515.45<br>p < 2e-16 (***) |
| (d) | 2.12<br>p = 0.977 | 1.40<br>p = 0.994 | 0.90<br>p = 0.999 | 0.23<br>p = 1.000 | 1.31<br>p = 0.995 |
| (e) | 15.91<br>p = 0.044 (*) | 5.37<br>p = 0.717 | 2.44<br>p = 0.964 | 5.41<br>p = 0.713 | 17.02<br>p = 0.030 (*) |
| (f) | 7.06<br>p = 0.530 | 4.53<br>p = 0.806 | 3.38<br>p = 0.908 | 5.37<br>p = 0.717 | 16.30<br>p = 0.038 (*) |
| (g) | 3.08<br>p = 0.929 | 1.68<br>p = 0.989 | 1.62<br>p = 0.990 | 2.90<br>p = 0.940 | 6.84<br>p = 0.554 |

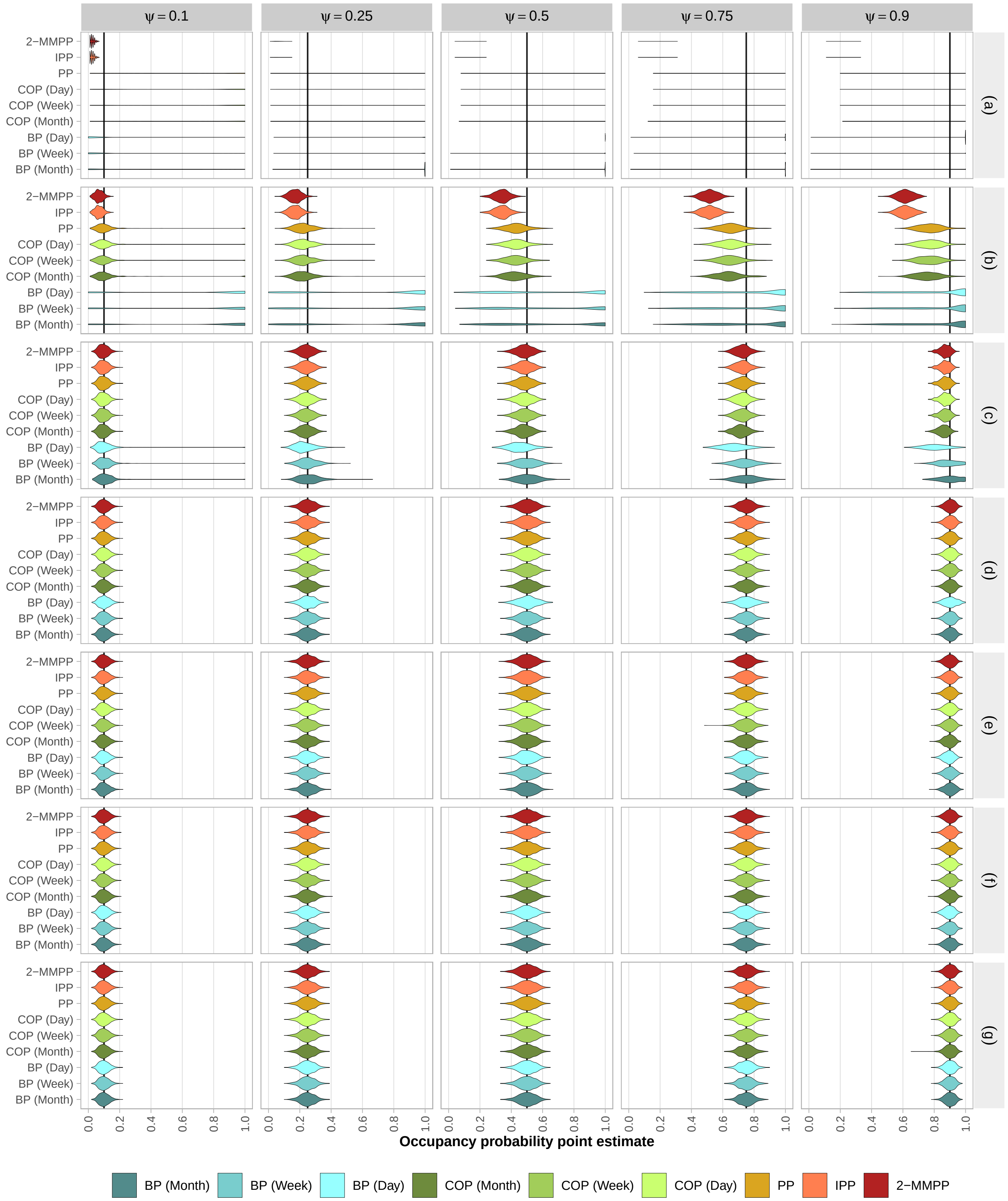


Figure S1: **Distribution of occupancy probability point-estimates ($\hat{\psi}$).** For each simulated scenario, with the simulated occupancy ($\psi$) on top and the simulated detectability (a to g, see Table 2) on the right. Models were fitted with the Nelder-Mead optimisation algorithm.

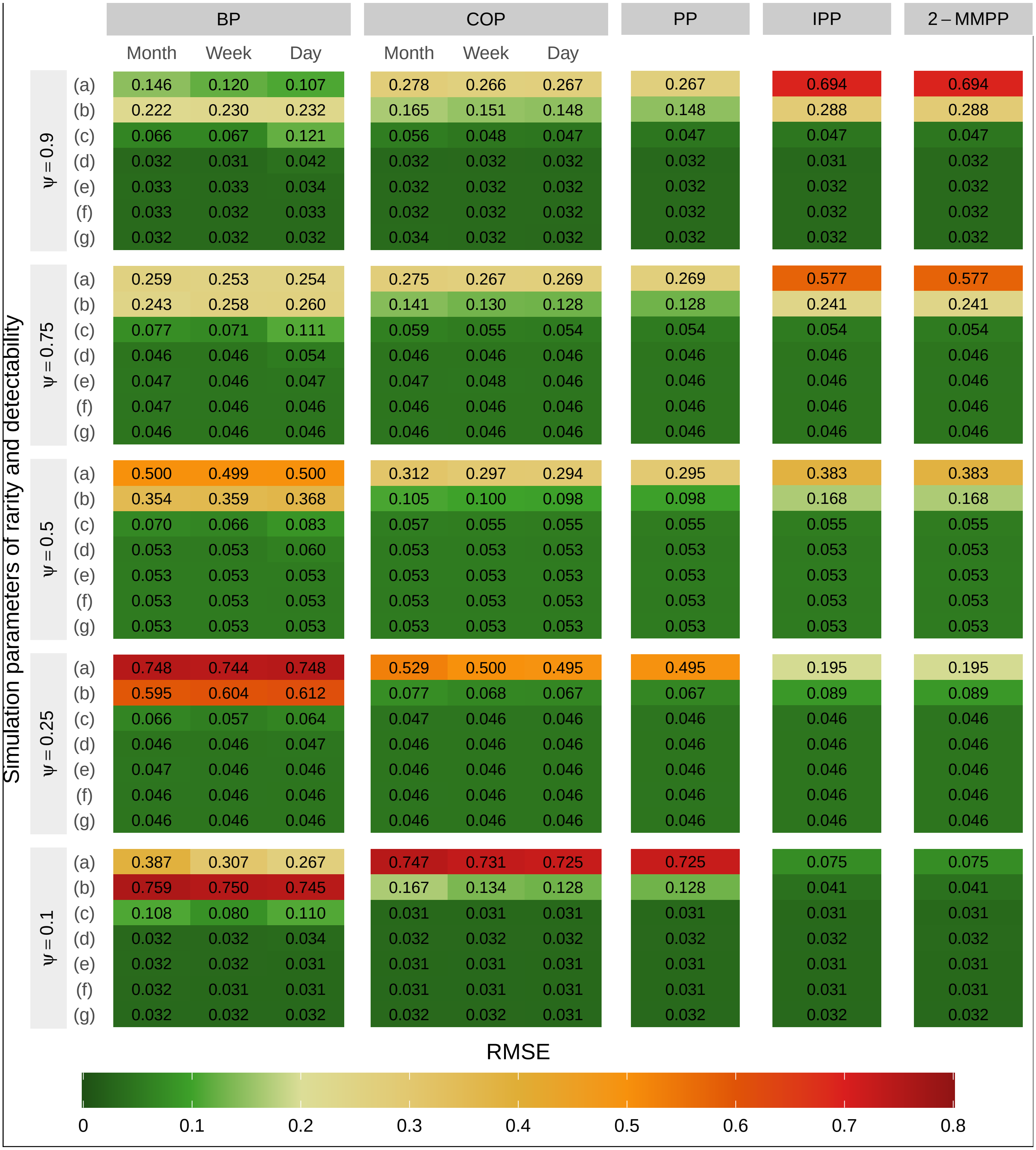


Figure S2: **Root Mean Squared Error (RMSE) of occupancy probability point-estimates ($\hat{\psi}$).** For each simulated scenario, with the simulated occupancy ($\psi$) on top and the simulated detectability (a to g, see Table 2). Models were fitted with the Nelder-Mead optimisation algorithm.

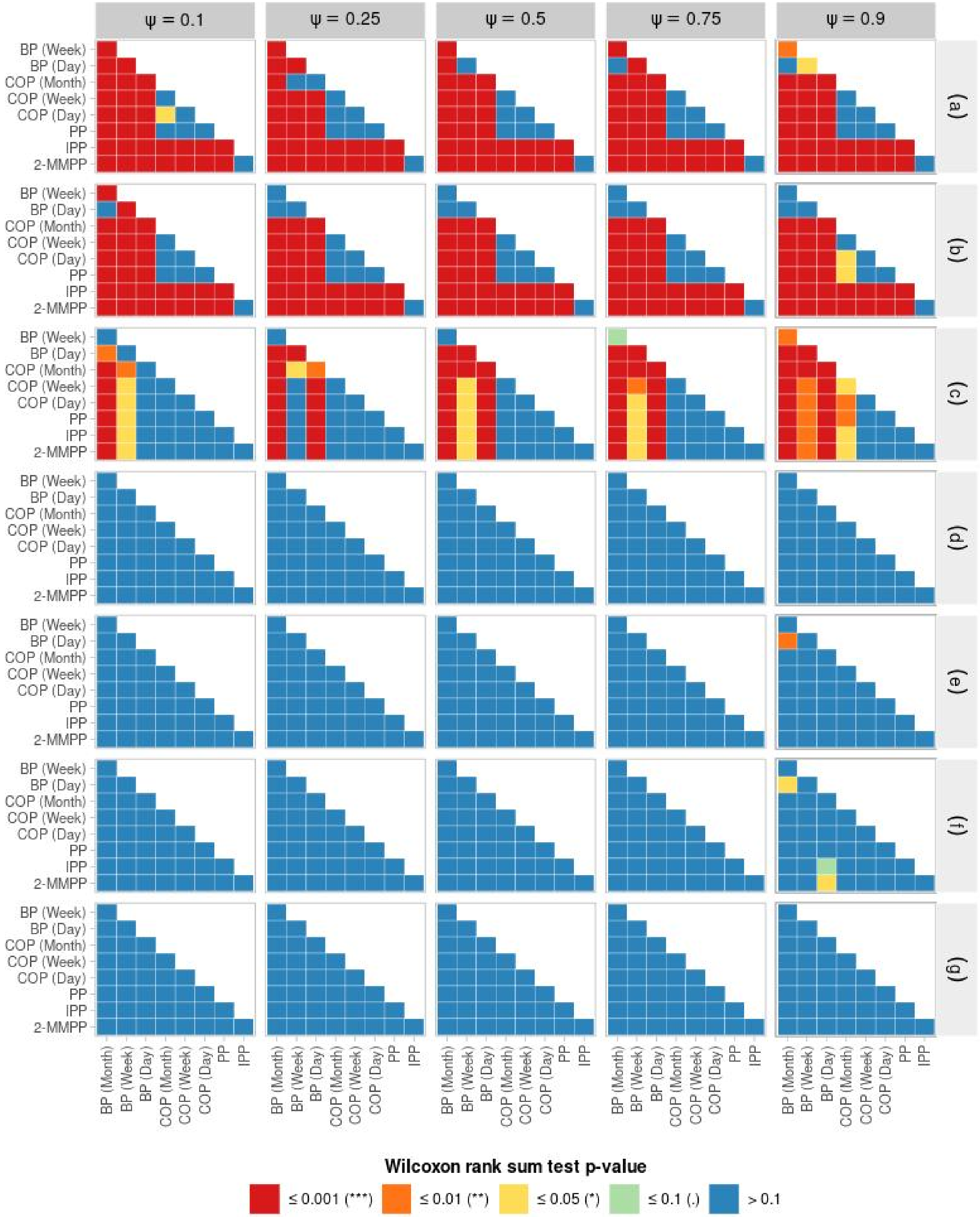


Figure S3: **Results of the Wilcoxon tests comparing the distribution of the occupancy probability point-estimates ($\hat{\psi}$) between models**. For each simulated scenario, with the simulated occupancy ($\psi$) on top and the simulated detectability (a to g, see Table 2) on the right. Models were fitted with the Nelder-Mead optimisation algorithm.

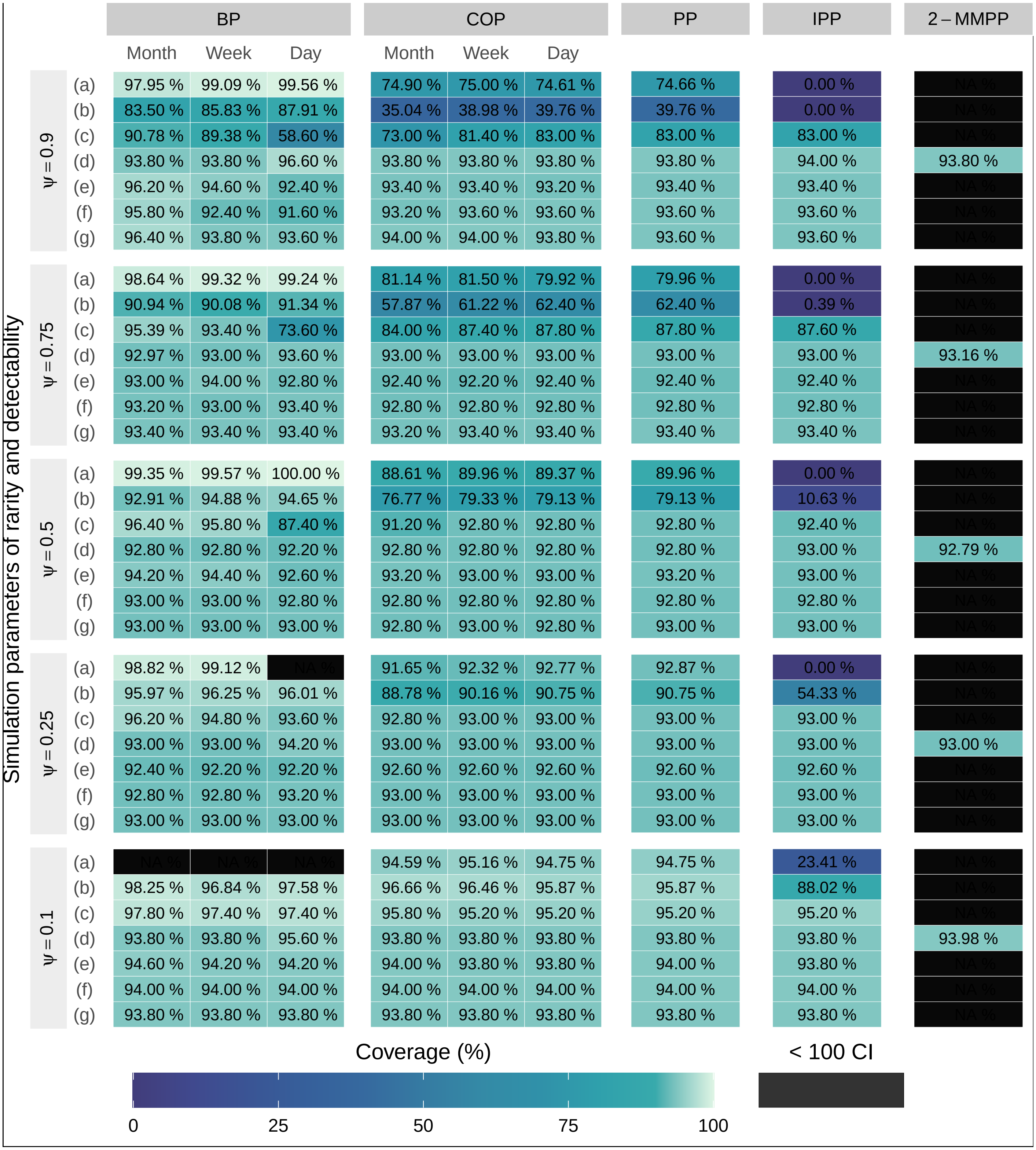


Figure S4: **Coverage, the proportion of simulations where the simulated value of $\psi$ is within the 95% confidence interval (CI) of $\hat{\psi}$.** For each simulated scenario, with the simulated occupancy ($\psi$) on top and the simulated detectability (a to g, see Table 2). Models were fitted with the Nelder-Mead optimisation algorithm. The coverage is not shown when there are less than 100 CI out of the 500 simulations that were calculated, because the Hessian matrix was not invertible.

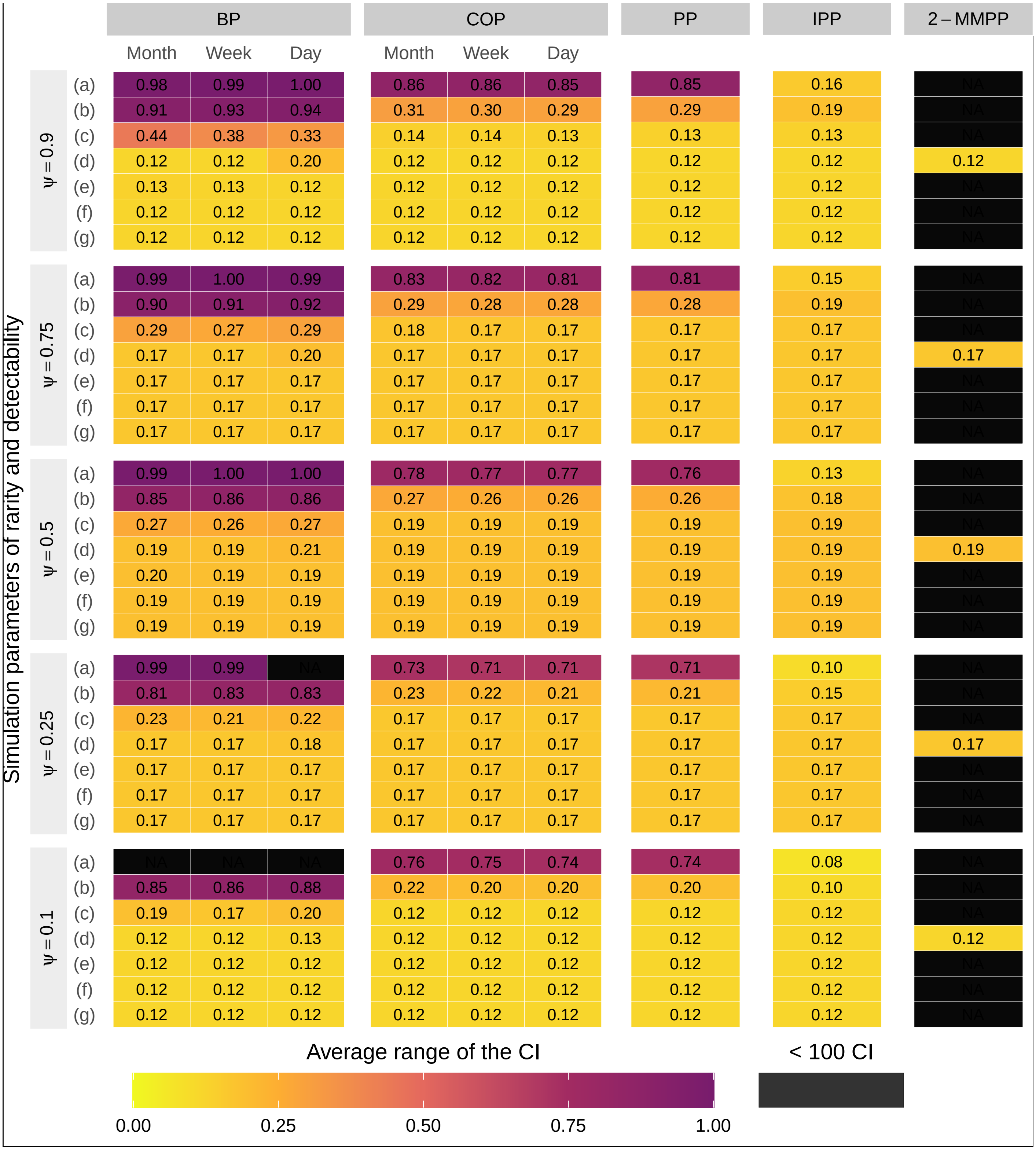


Figure S5: **Average range of the 95% confidence interval (ARCI) of $\hat{\psi}$.** For each simulated scenario, with the simulated occupancy ($\psi$) on top and the simulated detectability (a to g, see Table 2). Models were fitted with the Nelder-Mead optimisation algorithm. The ARCI is not shown when there are less than 100 CI out of the 500 simulations that were calculated, because the Hessian matrix was not invertible.

#### Appendix 5 Estimation with BFGS

In addition to maximizing likelihood using the Nelder-Mead optimization method, we explored the use of the BFGS algorithm, which is often the default option. We encountered more frequent optimisation errors with BFGS, attributed to non-finite differences, that could only be mitigated by adjusting the initial parameters. Consequently, the use of this algorithm produced incomplete results (Fig. S6, Fig. S7). With the BFGS algorithm, $\psi$ tended to be estimated more frequently at its boundaries, particularly in the BP model with daily sessions. Furthermore, the choice of initial parameters seemed to have a greater impact on $\psi$ estimation with BFGS compared to Nelder-Mead, indicating that BFGS may struggle more in reaching the maximum likelihood. Consequently, we recommend that model users exercise caution when selecting an optimisation algorithm. Further investigation could be undertaken to assess how the optimisation algorithm interacts with different models, determining whether certain optimisation algorithms are better suited depending on the model.

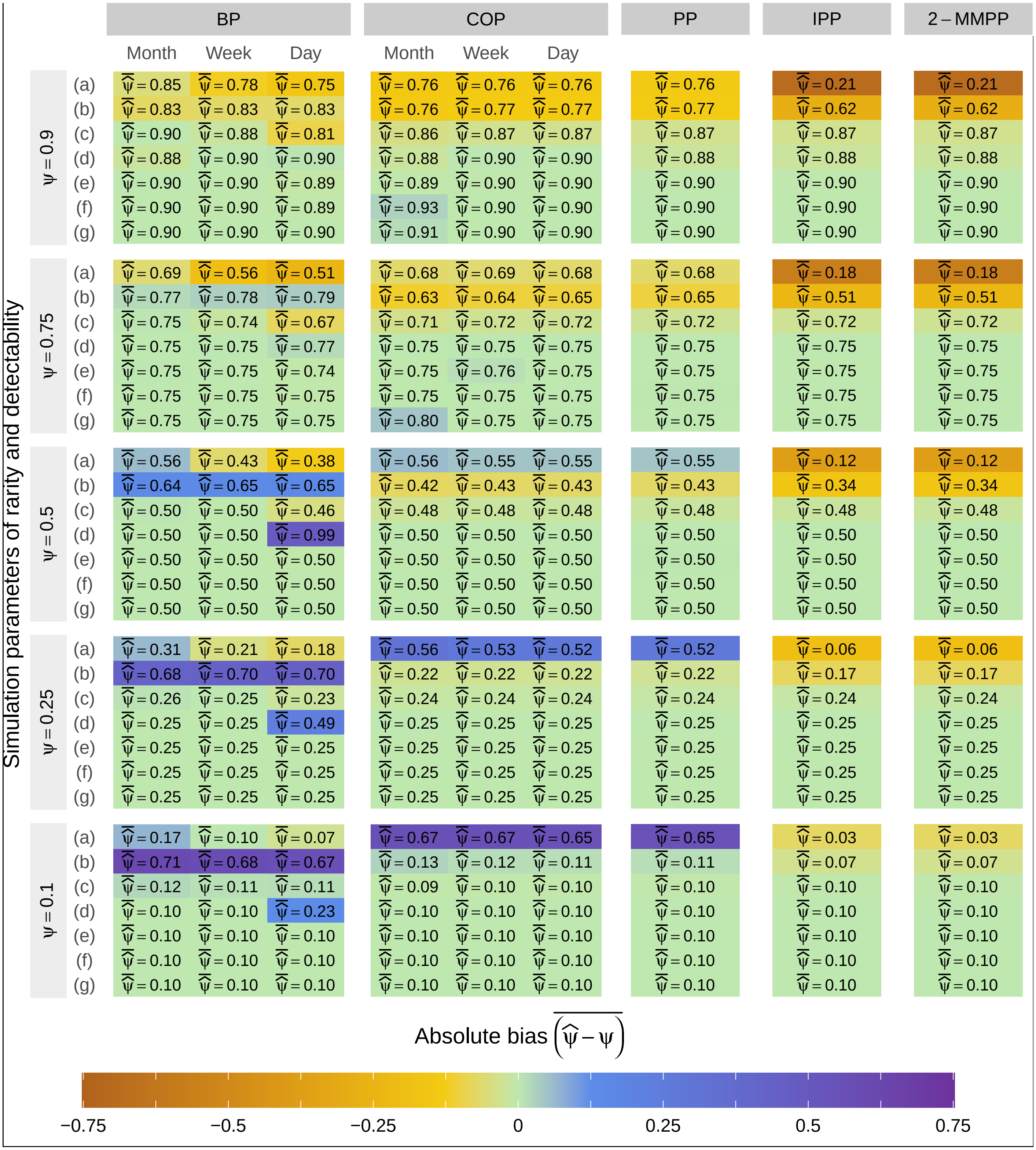


Figure S6: **Bias of occupancy probability point-estimates ($\hat{\psi}$).** For each simulated scenario, with the simulated occupancy ($\psi$) on top and the simulated detectability (a to g, see Table 2). Models were fitted with the **BFGS** optimisation algorithm.

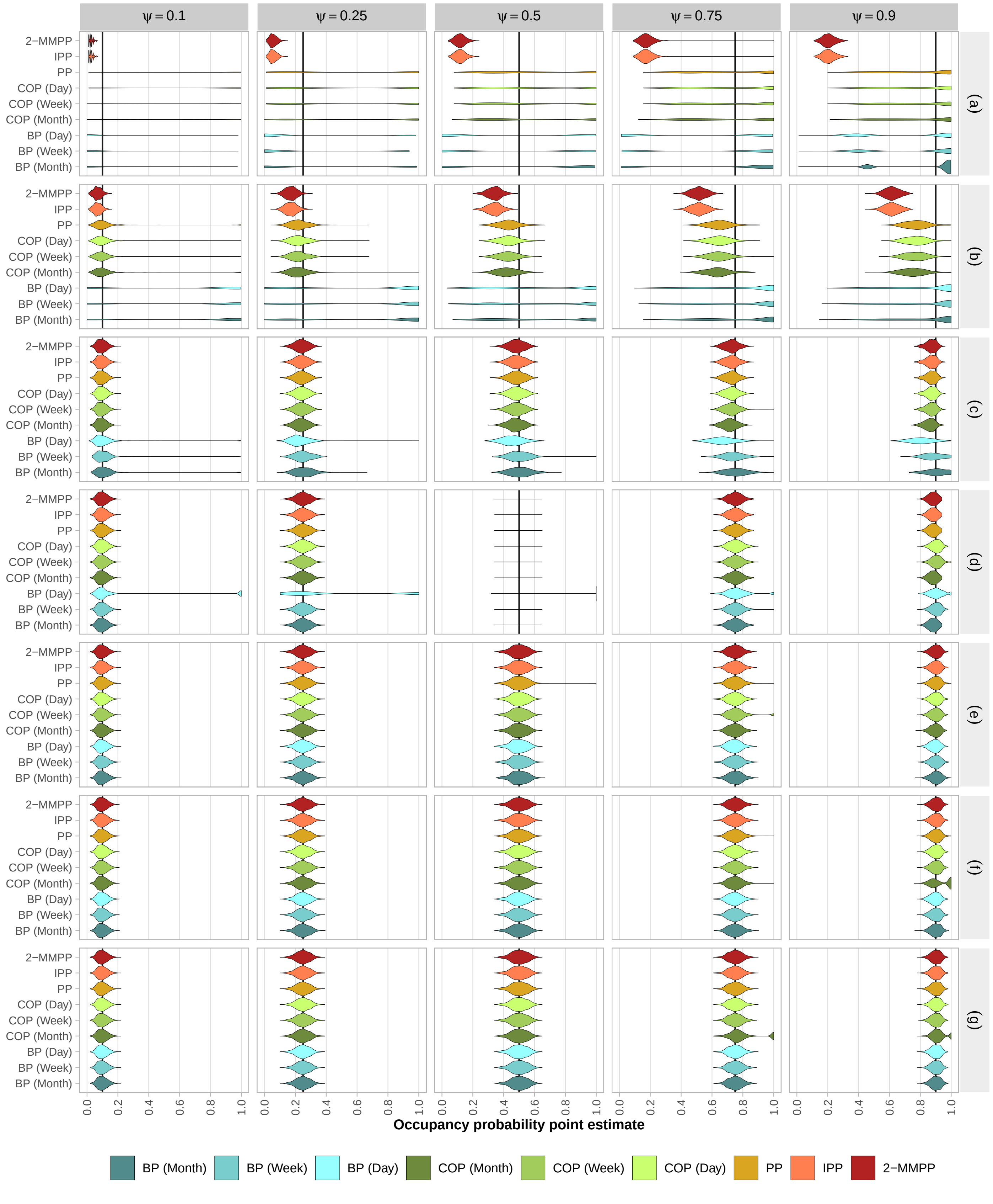


Figure S7: **Distribution of occupancy probability point-estimates ($\hat{\psi}$).** For each simulated scenario, with the simulated occupancy ($\psi$) on top and the simulated detectability (a to g, see Table 2) on the right. Models were fitted with the **BFGS** optimisation algorithm.

### Appendix 6 Application to lynx occupancy

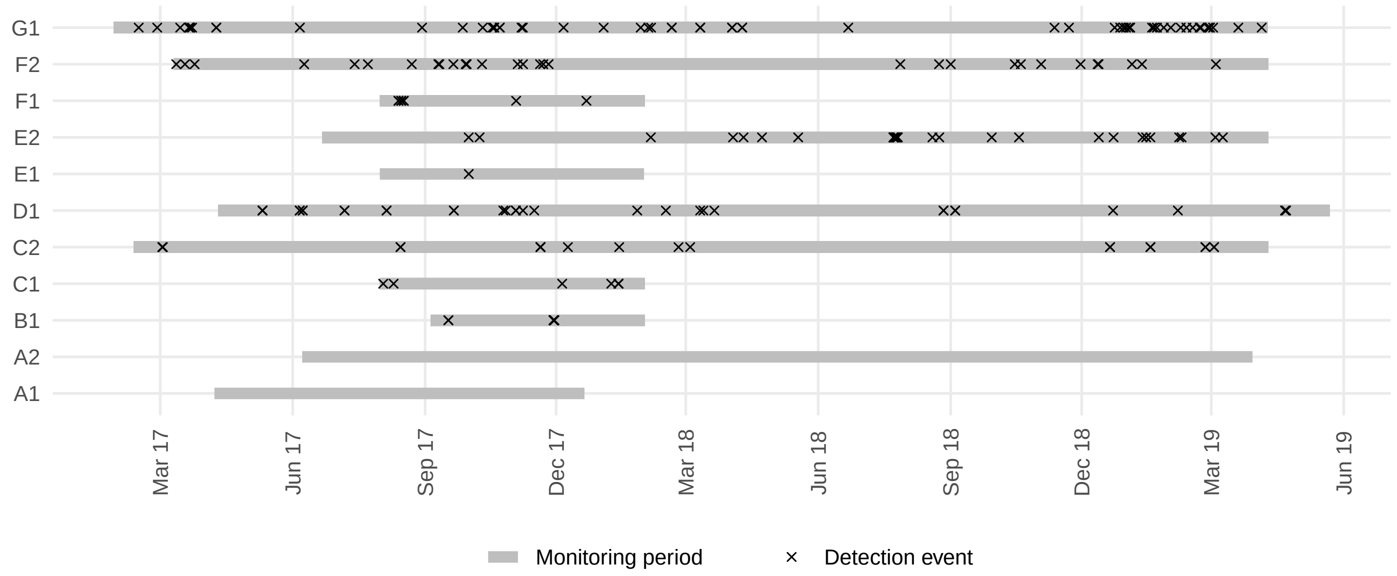


Figure S8: **Lynx continuous-time detection history**. Data collected by camera traps in 11 sites of the Ain county, France, from Gimenez et al. (2022).

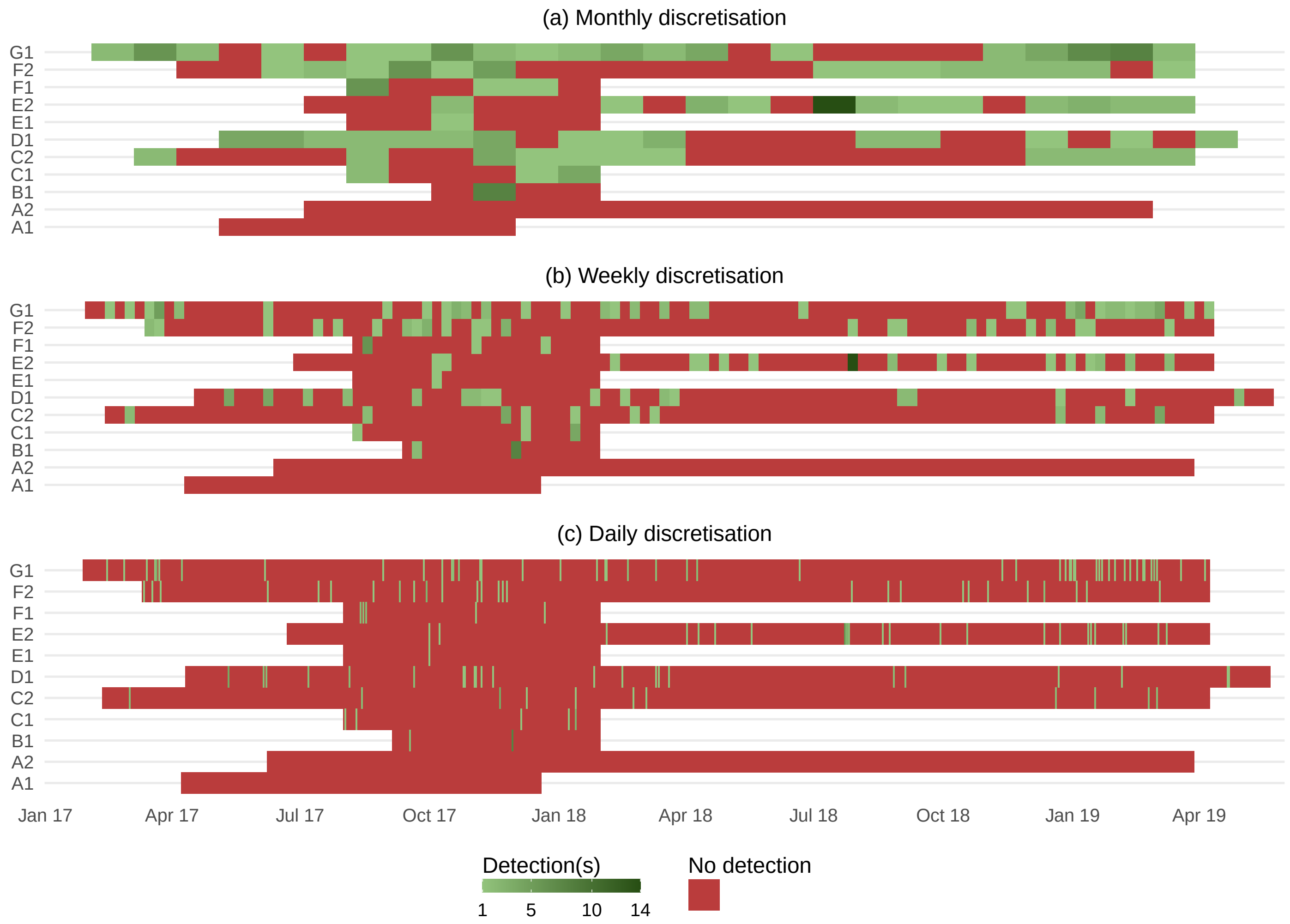


Figure S9: **Lynx discretised detection history**. Discretised in (a) monthly sessions, (b) weekly sessions, and (c) daily sessions.

Table S4: **Parameter estimates for all five models with the lynx data set**. For each model and discretisation (where relevant), we display the point estimates and the 95% and 50% confidence interval of each parameter. We also display the transformed parameter point estimate and standard error (SE). Probabilities ($\psi$, $p$) are logit-transformed and rates ($\lambda$, $\lambda_1$, $\lambda_2$, $\mu_{12}$, $\mu_{21}$) are log-transformed.

| Model | Sessions duration | Parameter | Point estimate | 95% CI | 50% CI | Transformed point estimate (± transformed SE) |
|---|---|---|---|---|---|---|
| BP | Month | $\psi$ | 0.751 | 0.414 - 0.928 | 0.647 - 0.833 | 1.106 (± 0.742) |
| | | $p$ | 0.370 | 0.292 - 0.456 | 0.342 - 0.399 | -0.531 (± 0.181) |
| BP | Week | $\psi$ | 0.767 | 0.414 - 0.939 | 0.66 - 0.849 | 1.193 (± 0.786) |
| | | $p$ | 0.079 | 0.06 - 0.104 | 0.072 - 0.087 | -2.453 (± 0.153) |
| BP | Day | $\psi$ | 0.800 | 0.41 - 0.959 | 0.687 - 0.88 | 1.389 (± 0.895) |
| | | $p$ | 0.008 | 0.006 - 0.012 | 0.007 - 0.009 | -4.785 (± 0.172) |
| COP | Month | $\psi$ | 0.818 | 0.493 - 0.954 | 0.727 - 0.884 | 1.504 (± 0.782) |
| | | $\lambda$ | 0.046 | 0.04 - 0.052 | 0.042 - 0.046 | -3.089 (± 0.071) |
| COP | Week | $\psi$ | 0.818 | 0.493 - 0.954 | 0.726 - 0.884 | 1.504 (± 0.782) |
| | | $\lambda$ | 0.046 | 0.04 - 0.052 | 0.042 - 0.046 | -3.085 (± 0.07) |
| COP | Day | $\psi$ | 0.818 | 0.493 - 0.954 | 0.726 - 0.884 | 1.504 (± 0.782) |
| | | $\lambda$ | 0.045 | 0.04 - 0.052 | 0.042 - 0.045 | -3.092 (± 0.07) |
| PP | | $\psi$ | 0.818 | 0.493 - 0.954 | 0.727 - 0.884 | 1.506 (± 0.782) |
| | | $\lambda$ | 0.045 | 0.039 - 0.052 | 0.041 - 0.045 | -3.094 (± 0.07) |
| 2-MMPP | | $\psi$ | 0.818 | 0.493 - 0.954 | 0.727 - 0.884 | 1.504 (± 0.782) |
| | | $\lambda_1$ | 0.008 | 0.001 - 0.059 | 0.004 - 0.016 | -4.783 (± 0.996) |
| | | $\lambda_2$ | 17.644 | 9.435 - 32.995 | 14.225 - 21.885 | 2.87 (± 0.319) |
| | | $\mu_{12}$ | 0.101 | 0.035 - 0.294 | 0.07 - 0.146 | -2.291 (± 0.545) |
| | | $\mu_{21}$ | 48.185 | 32.331 - 71.813 | 42.003 - 55.277 | 3.875 (± 0.204) |
| IPP | | $\psi$ | 0.818 | 0.493 - 0.954 | 0.726 - 0.884 | 1.502 (± 0.781) |
| | | $\lambda_2$ | 14.221 | 9.237 - 21.895 | 12.259 - 16.498 | 2.655 (± 0.22) |
| | | $\mu_{12}$ | 0.162 | 0.119 - 0.221 | 0.146 - 0.18 | -1.82 (± 0.158) |
| | | $\mu_{21}$ | 50.704 | 34.608 - 74.287 | 44.459 - 57.826 | 3.926 (± 0.195) |
